# Focal Laser Stimulation of Fly Nociceptors Activates Distinct Axonal and Dendritic Ca^2+^ Signals

**DOI:** 10.1101/2021.03.04.433797

**Authors:** Rajshekhar Basak, Sabyasachi Sutradhar, Jonathon Howard

## Abstract

*Drosophila* Class IV neurons are polymodal nociceptors that detect noxious mechanical, thermal, optical and chemical stimuli. Escape behaviors in response to attacks by parasitoid wasps are dependent on Class IV cells, whose highly branched dendritic arbors form a fine meshwork that is thought to enable detection of the wasp’s needle-like ovipositor barb. To understand how mechanical stimuli trigger cellular responses, we used a focused 405-nm laser to create highly local lesions to probe the precise position needed in evoke responses. By imaging calcium signals in dendrites, axons, and soma in response to stimuli of varying positions, intensities and spatial profiles, we discovered that there are two distinct nociceptive pathways. Direct stimulation to dendrites (the contact pathway) produces calcium responses in axons, dendrites and the cell body whereas stimulation adjacent to the dendrite (the non-contact pathway) produces calcium responses in the axons only. We interpret the non-contact pathway as damage to adjacent cells releasing diffusible molecules that act on the dendrites. Axonal responses have higher sensitivities and shorter latencies. In contrast, dendritic responses have lower sensitivities and longer latencies. Stimulation of finer, distal dendrites leads to smaller responses than stimulation of coarser, proximal dendrites, as expected if the contact response depends on the geometric overlap of the laser profile and the dendrite diameter. Because the axon signals to the CNS to trigger escape behaviors, we propose that the density of the dendritic meshwork is high not only to enable direct contact with the ovipositor, but also to enable neuronal activation via diffusing signals from damaged surrounding cells. Dendritic contact evokes responses throughout the dendritic arbor, even to regions distant and distal from the stimulus. These dendrite-wide calcium signals may facilitate hyperalgesia or cellular morphological changes following dendritic damage.

**Statement of Significance:** Animals encounter a wide range of noxious stimuli in the natural world. Nociceptive neurons are specialized cells that sense harmful stimuli and trigger avoidance responses. Class IV cells, located under the cuticle in *Drosophila* larvae, are polymodal nociceptors that respond to noxious mechanical, thermal, optical, and chemical stimuli. To investigate the spatial requirements of mechanoreception in Class IV neurons, we measured calcium signals evoked by a focused laser beam that creates highly localized tissue damage. We discovered that different cellular compartments – axons and dendrites – responded differentially depending on whether the stimulus makes direct contact with the neuron or not. This provides evidence that mechanical nociception in Class IV cells occurs via two distinct pathways.

## Introduction

Nociception is the sensation of painful or injurious stimulation. The peripheral nervous system senses noxious stimuli through nociceptive cells, which signal to the central nervous systems to trigger appropriate behavioral responses (Fields, 1987; Tracey, 2017). While much is known about the molecular basis of thermal, chemical, and mechanical nociception (Basbaum et al., 2009; McCleskey and Gold, 1999), many questions remain. For example, injurious mechanical stimuli are difficult to replicate reliably and could have multiple direct effects the nociceptors, or may have indirect effects via damage to cells in the surrounding tissue. Therefore, elucidating the transduction pathways for nociceptive mechanical stimuli is likely to be difficult.

*Drosophila* Class IV dendritic arborization (da) neurons are polymodal nociceptors that serve as a model system for studying nociception (Im and Galko, 2012). These highly branched cells (Grueber et al., 2002) innervate the epidermis of the larval body and respond to noxious mechanical (Guo et al., 2014; Kim et al., 2012; Zhong et al., 2010), thermal (Babcock et al., 2009; Terada et al., 2016; Tracey et al., 2003), chemical (Lopez-Bellido et al., 2019) and optical (Xiang et al., 2010; Yamanaka et al., 2013) stimuli. Noxious stimulation triggers avoidance behaviors in the larvae that are attenuated when these cells are specifically ablated (Tracey et al., 2003). A striking ecological example of nociception by this cell is the larval avoidance response to attacks by parasitoid wasps, which puncture the cuticle with their ovipositors to lay eggs in the larvae (Hwang et al., 2007; Robertson et al., 2013). Silencing Class IV neurons alone resulted in loss of defensive rolling escape behaviors (Hwang et al., 2007). The dense network of Class IV dendrites, which have a mesh size of several microns (Ganguly et al., 2016), may increase the likelihood that an ovipositor, which has a diameter that tapers from 20 μm down to 1 μm (Robertson et al., 2013), makes direct contact with the arbor.

The question we address is how the ovipositor stimulates the Class IV cell. It is reported that penetration of the larval cuticle by the wasp’s ovipositor can physically damage the fine dendrites of Class IV cells (Tracey, 2017). If the damage directly punctures the dendrite’s plasma membrane, this could lead to a local depolarization of the membrane potential, which could propagate electrotonically or by action potentials to the cell body and axon, and then to the central nervous system to trigger an escape behavior. Alternatively, it is possible that the ovipositor damages other cells such as epidermal or muscle cells, which in turn signal to the Class IV cells via factors released into the extracellular medium or through acidification; these factors or protons could then bind to receptors on the membrane of Class IV dendrites, leading ultimately to the opening of ion channels and receptors potentials that propagate to the cell body and axon (Tracey, 2017). This indirect pathway would be analogous to the P2X3 receptors of vertebrate nociceptors that bind to ATP released by damaged cells (Cook and McCleskey, 2002; Hamilton and McMahon, 2000).

In this study, we investigated potential direct and indirect nociceptive mechanisms using a focused laser beam to locally damage Class IV neurons and/or the adjacent tissue. We then used the genetically encoded calcium reporter GCaMP6f (Chen et al., 2013) to test whether the stimuli evoked calcium responses in Class IV cells. We found that stimuli trigger two distinct calcium signaling responses based on their location with respect to the Class IV dendrites. If the laser damages the adjacent tissue but not the dendrite, then robust calcium responses are recorded in the axons, but not in the dendrites or cell bodies. If the laser damages the dendrites, then robust calcium responses are recorded throughout the dendrite and cell body, in addition to the axon. Thus, Class IV cells are excited by both direct and indirect nociceptive pathways.

## Results

### Focal 405-nm stimulation triggers cuticular damage, behavioral responses and intracellular calcium increases

To study nociception by Class IV neurons, we used a focused 405-nm laser beam to mimic penetration of the larval cuticle by the wasp ovipositor. We irradiated individual Class IV neurons in unanesthetized larvae that had been constrained in a PDMS device (Mishra et al., 2014) mounted on the stage of a spinning disk confocal microscope (Figure 1A). When focused to a diameter of 0.5 or 1 μm (full width at half maximum, FWHM), laser illumination with integrated power ≥80% (≥32 mW, *Supplementary Figure* 1) and duration 0.2 s produced cuticular puncture wounds (Fig 1B), severed dendrites, and caused bleaching that did not recover over 20 minutes (*Supplementary Figure* 2A,B). The diameters of the puncture wounds were 2-4 μm (Figure 1C), similar to the diameters of wasp ovipositor barbs, though smaller than the maximum 20-μm diameter of the ovipositor itself (Robertson et al., 2013). These laser powers produced “melanotic spots” (inset to Figure 1B), a characteristic of cuticular penetration by the ovipositor (Galko and Krasnow, 2004; Robertson et al., 2013). Integrated powers ≤40% did not produce punctures; at these intensities bleaching of illuminated dendrites occurred, but recovered over 20 minutes (*Supplementary Figure* 2C,D).

**Figure 1.**
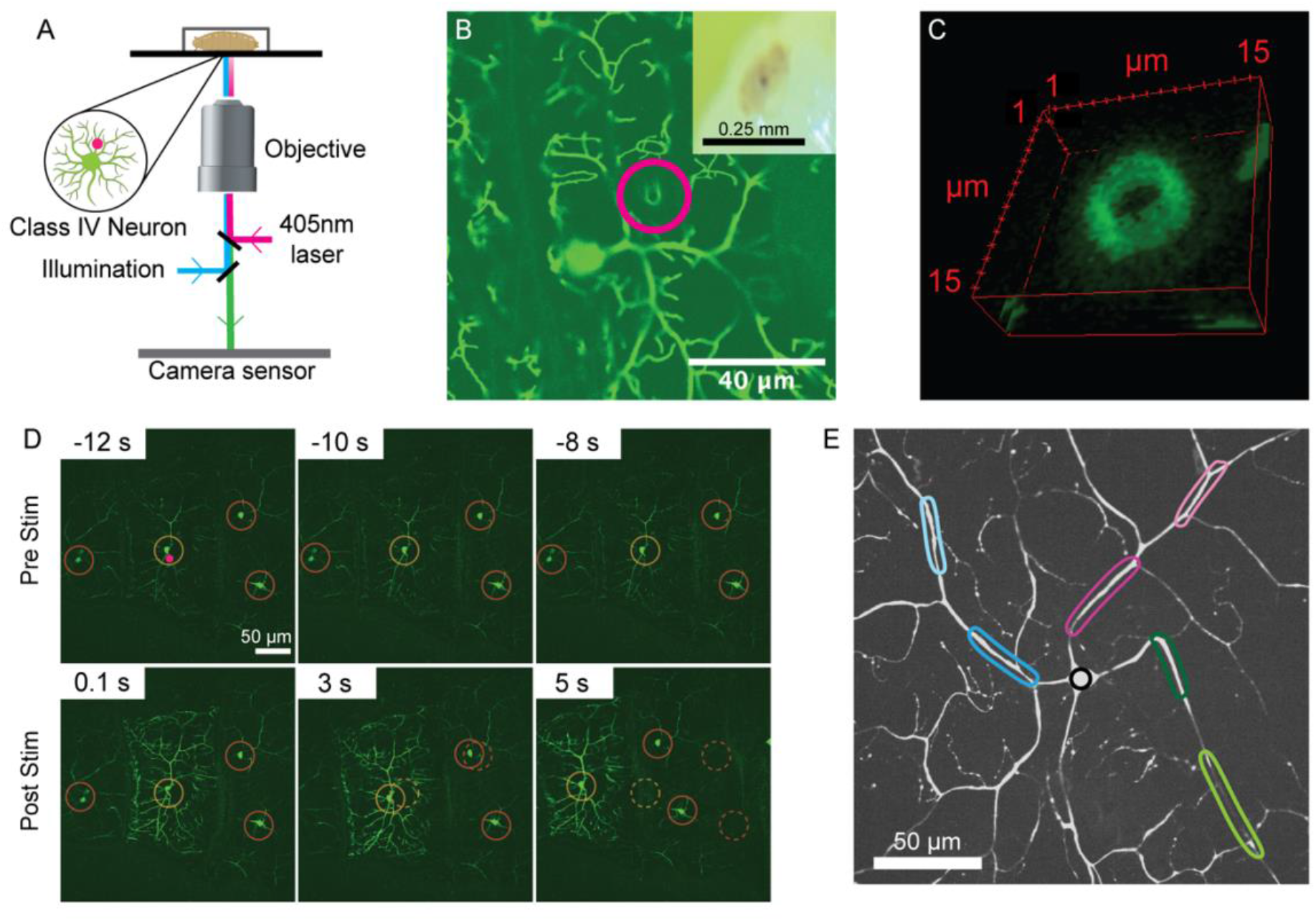
Focused 405-nm laser stimulation triggers larval behavioral responses and calcium signals in Class IV cells. (A) Schematic diagram depicting the stimulation and imaging setup. (B) Example of a puncture wound (magenta circle) in the dendritic arbor of a Class IV neuron expressing GFP. Inset shows a melanotic spot at the site of illumination. (C) Z-stack slices of puncture wound. (D) Montage depicting behavioral and calcium responses to 405-nm stimulation at 80% stimulation intensity. Top row shows larva stationary before stimulation; bottom row shows tissue movement and calcium increase measured using the genetically encoded calcium reporter GCaMP6f. Dashed circles indicate positions of the cell bodies prior to stimulation. (E) Image of a Class IV neuron depicting seven regions of interest (ROI): four on dendrites (magenta and blue), two on the axon (green) and one on the soma (black). Darker color is more proximal.

Constrained larvae pulsed with the 405-nm laser at ≥80% power for 0.1 s exhibited behavioral responses that manifest as tissue movements (Figure 1D) (*Supplementary Movie 1-2*). Unconstrained larvae writhed, crawled, and turned upon laser irradiation, with higher stimulation intensity eliciting a stronger response. The laser stimulation was not lethal: all six larvae (70 hrs AEL) subject to ≥80% maximum power survived for 24 hours.

Focal laser stimulation within the dendritic fields of Class IV neurons induced intracellular calcium increases. Following pulsed stimulation for 0.1 s at 80% power, the fluorescence of the calcium indicator GCaMP6f, expressed specifically in Class IV cells (see Methods), increased (Fig 1D). The fluorescence increases could be observed in the cell body, the dendrites and the axon, with amplitudes up to several-fold above baseline and lasting for several seconds (*Supplementary Movie 1-2*). The fluorescence change was mediated, at least in part, by calcium influx through voltage-gated calcium channels: RNAi of the Ca-alpha1D sub-unit of voltage-gated calcium channels in Class IV neurons resulted in smaller fluorescence changes (*Supplementary Figure 5*), as has been found for thermal responses in these cells (Terada et al., 2016). Thus, our focused 405-nm laser stimulus is a non-lethal nociceptive stimulus that mimics cuticle penetration by an ovipositor barb, producing both behavioral and cellular responses. The laser stimulus has advantages over an attack by a wasp’s ovipositor, as its position, intensity, geometry and duration can be controlled precisely.

### The calcium response depends on the position of the 405-nm illumination

To test whether physical damage to Class IV dendrites is necessary for nociceptive responses, we took advantage of the narrow spatial profile of our laser probe, as well as our ability to precisely control its position relative to the dendritic processes. We found that larval behavioral responses were triggered irrespective of the stimulus location. However, we observed different calcium responses in Class IV neurons depending on whether the stimulus made direct contact with the dendritic arbor or not. The background fluorescence of the GCaMP6f-expressing cells was sufficiently high to unambiguously identify even distal dendritic processes (e.g. Figure 1E). We found that non-contact illumination generated calcium transients in the axons of Class IV neurons, but greatly attenuated signals in the dendrites (Figure 2A,B; *Supplementary Movie 3*). In contrast, direct contact of the stimulus with the arbor results in calcium responses throughout the entire cell (Figure 2C,D; *Supplementary Movie 4*). To confirm this result, we repeated the same experiment by stimulating single cells three times in three different places: the first two stimuli made no contact with Class IV arbors while the third made contact. We found that only axons responded when no contact was made, whereas axon, dendrite and soma all responded when contact was made (Fig 2E). Thus, there are two distinct calcium signaling responses: (i) a “non-contact” response in axons only, and (ii) a “contact” response in all compartments.

**Fig 2.**
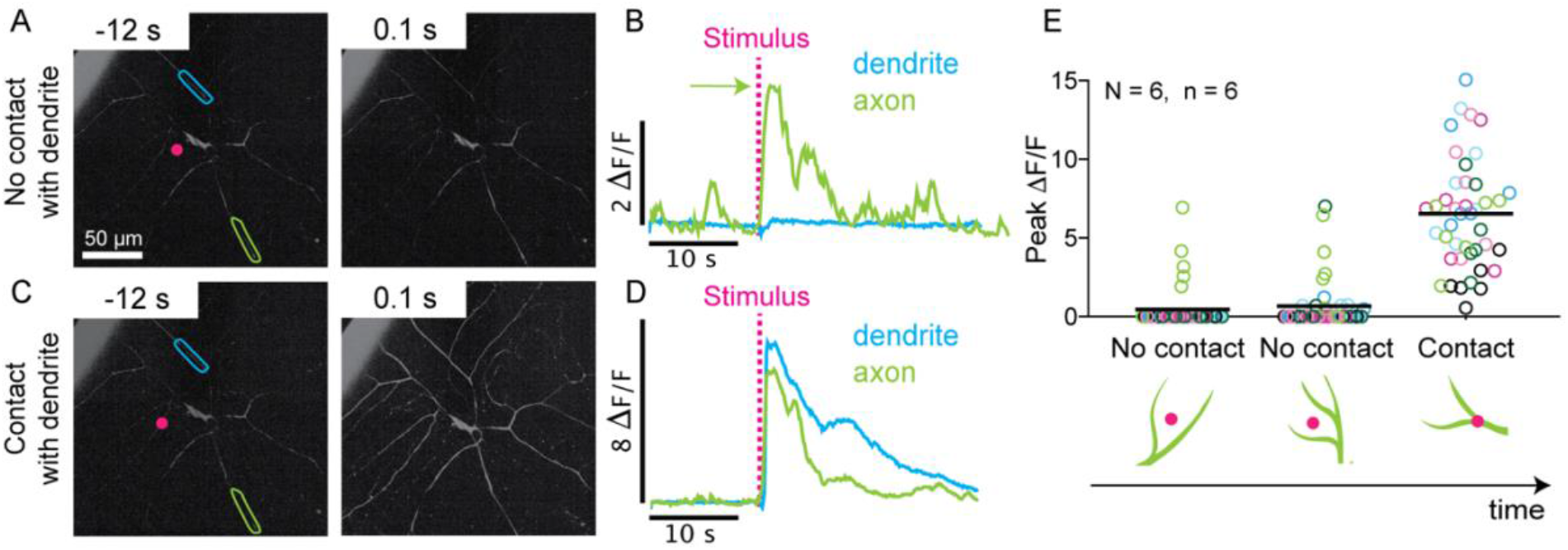
405-nm stimulation triggers two distinct calcium signaling responses in Class IV neurons. (A, B) When the laser is focused adjacent to but not on a dendrite (non-contact), large calcium changes are recorded in the axon (green ROI) and only very small changes are recorded in the dendrite (blue ROI). (C, D) When the laser is focused on a dendrite (contact), large calcium changes are recorded in both the axon (green ROI) and dendrite (blue ROI). (E) Magnitude of normalized fluorescence responses across all 7 ROIs (open circles color coded as in Figure 1E) for a cell stimulated 3 consecutive times at different locations. The first two stimulations did not make contact with dendritic arbor and evoked axonal responses only (green); the third stimulation made contact and evoked responses everywhere. Black lines indicate means for all ROIs combined. N represents number of larvae; n represents number of cells.

### The non-contact response

To investigate the conditions under which the non-contact response in axons is triggered, we probed larvae with laser stimuli of different intensities (up to 100% laser power of 45mW) and spatial profiles (full width at half maximum equal to 0.5 or 1 μm; Figure 3A). We first asked how the likelihood of a calcium response depended on the light intensity. A region of interest (ROI, defined in Figure 1E) was deemed responsive if the relative change in fluorescence, *ΔF*/*F*, after stimulation was larger than five standard deviations of the baseline fluorescence, *F*, prior to stimulation (*Materials and Methods*). We found that the percentage of axonal ROIs that responded to non-contact stimuli increased from 10% (FWHM 1 μm) and 30% (FWHM 0.5 μm) at 10% laser power to 70-80% at 100% laser power (Figure 3B). Thus, the non-contact stimulus reproducibly evokes responses from Class IV axons, with narrower profiles giving larger responses at lower total intensities.

**Fig 3.**
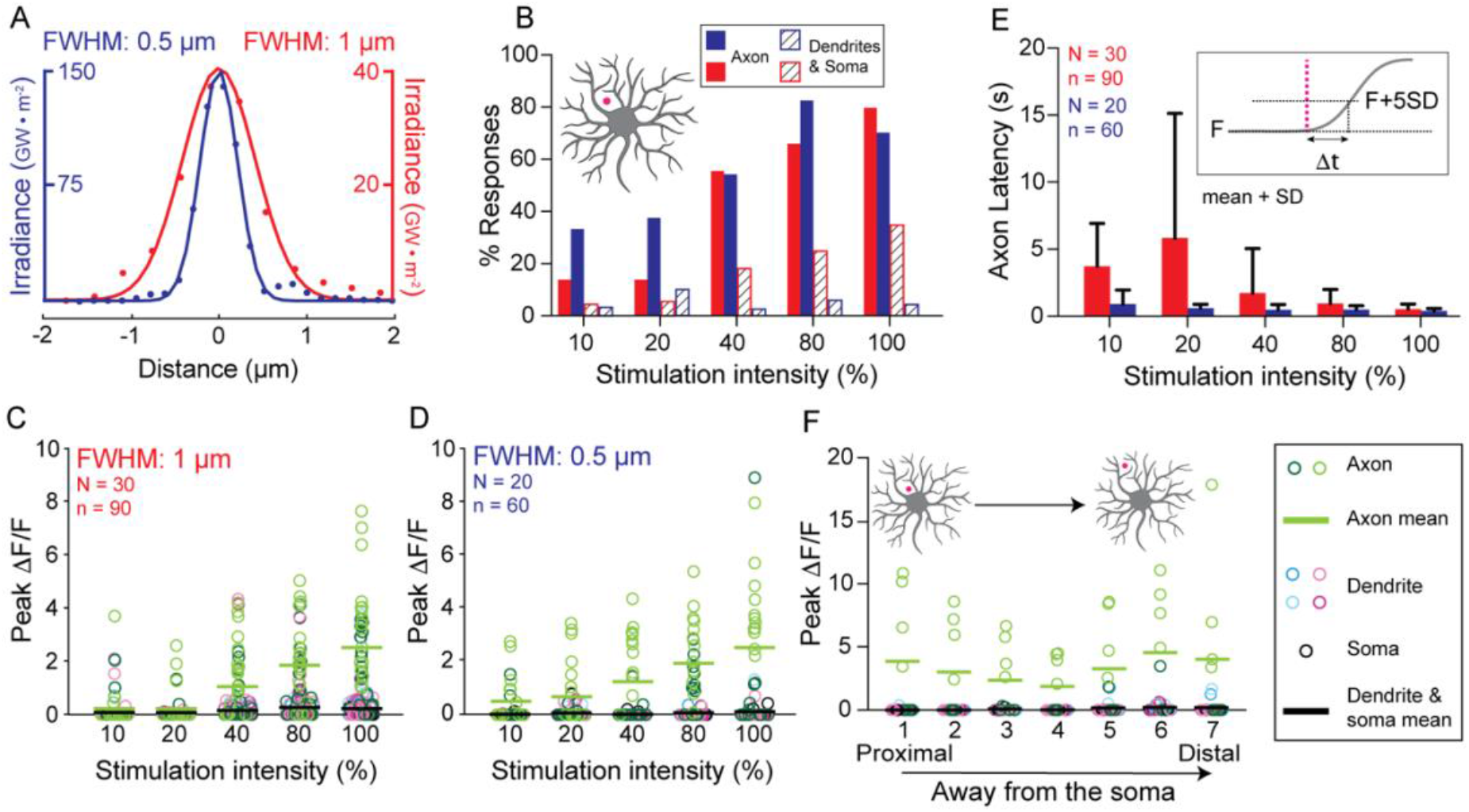
Characterization of the “non-contact” axonal calcium response. (A) Line scans of the spatial profiles of the narrower 405-nm profile (0.5 μm FWHM, blue) and the wider profile (1 μm FWHM, red). When they have the same total power (intensity), the irradiance (on the *y*-axes) of the narrower profile is four times larger. (B) Frequency of calcium transients in the two axonal ROIs (solid bars) and the five dendritic and somal ROIs (striped bars) in response to non-contact stimulation across a range of intensities (10 - 100%). Red and blue correspond to wider and narrower profiles. (C-D) Peak *ΔF*/*F* values for cells stimulated with no contact. Open circles indicate ROIs color-coded as in Fig 1E. Lines denote means of axon ROIs (green) and dendrite/soma ROIs (black). Statistical comparisons for these data are in *Supplementary Table 1*. (E) Axonal response latencies. Red and blue histograms correspond to wider and narrower stimuli. Inset: the latency is defined as the time when *ΔF*/*F* = *F* + 5SD. Statistical comparisons for data are shown in *Supplementary Table 2*. (F) Peak values of *ΔF*/*F* for seven consecutive non-contact stimuli (0.5 μm FWHM) at increasing distances from the cell body. Ordinary One-way ANOVA test shows no difference between axon means (*p* = 0.8501).

We found that the magnitudes of calcium responses were also graded with stimulation intensity. The peak value of *ΔF*/*F* in the axon following the laser pulse increased from an average of 0.2 (FWHM 1 μm) and 0.5 (FWHM 0.5 μm) at 10% laser paper to an average of ∼2.5 at 100% laser power for both stimulation profiles (Figure 3C,D). Thus, axonal calcium transients induced by non-contact stimulation are not all-or-nothing but rather graded with stimulus intensity.

In contrast to the axonal responses, only a small fraction of dendritic and somal ROIs responded to non-contact stimulation (Fig 3B, striped bars). Furthermore, the magnitudes of these responses were small: the average *ΔF*/*F*, was ≪ 1, with few ROIs giving *ΔF*/*F* > 0.1 even at the highest intensities (Figure 3C, D, black lines).

The latencies of the non-contact axonal responses, defined in the inset to Figure 3E, decreased with increasing intensity (Figure 3E). The narrower stimulus (0.5 μm FWHM) gave shorter latencies than the wider stimulus. For example, the latencies at 100% power were 0.39 ± 0.17 s (mean ± SD, n=12) for 0.5 μm FWHM and 0.52 ± 0.38 s (mean ± SD, n=18) for 1 μm FWHM.

The non-contact axonal response did not depend on the proximity of the stimulus to the cell body or the axon. To test this, we stimulated cells seven times at 80% intensity, with each stimulus progressively further away from the soma. We found no consistent effect of stimulus location (Figure 3F) (ordinary one-way ANOVA, *p* = 0.8501 was not significant).

In summary, the non-contact axonal response is graded, with higher intensities leading to a higher likelihood of responding, a larger fluorescence change when responding, and a shorter latency. By contrast, dendrites and soma responded infrequently to non-contact stimulation and the responses were much smaller.

### The contact response

To investigate the conditions under which the contact response is triggered, we illuminated the Class IV cells directly with the focused laser in three different locations: soma (Figure 4A), proximal dendrites (Figure 4B) and distal dendrites (Figure 4C). To quantify the likelihood of responses for each stimulation condition, we computed the percentage of the two axonal ROIs and the five dendrite and soma ROIs that responded to stimuli of different intensities and spatial profiles. We found that direct stimulation of the soma, proximal dendrites, and distal dendrites all evoked calcium transients throughout the cell with the percentage responding increasing with increasing stimulus intensity (Figure 4A-C). At the highest intensities, most ROIs responded, with the soma and proximal stimulation being somewhat more efficacious than distal illumination. In contrast to the non-contact response, the percentage of contact responses evoking calcium responses was similar in dendrites and axons.

**Figure 4.**
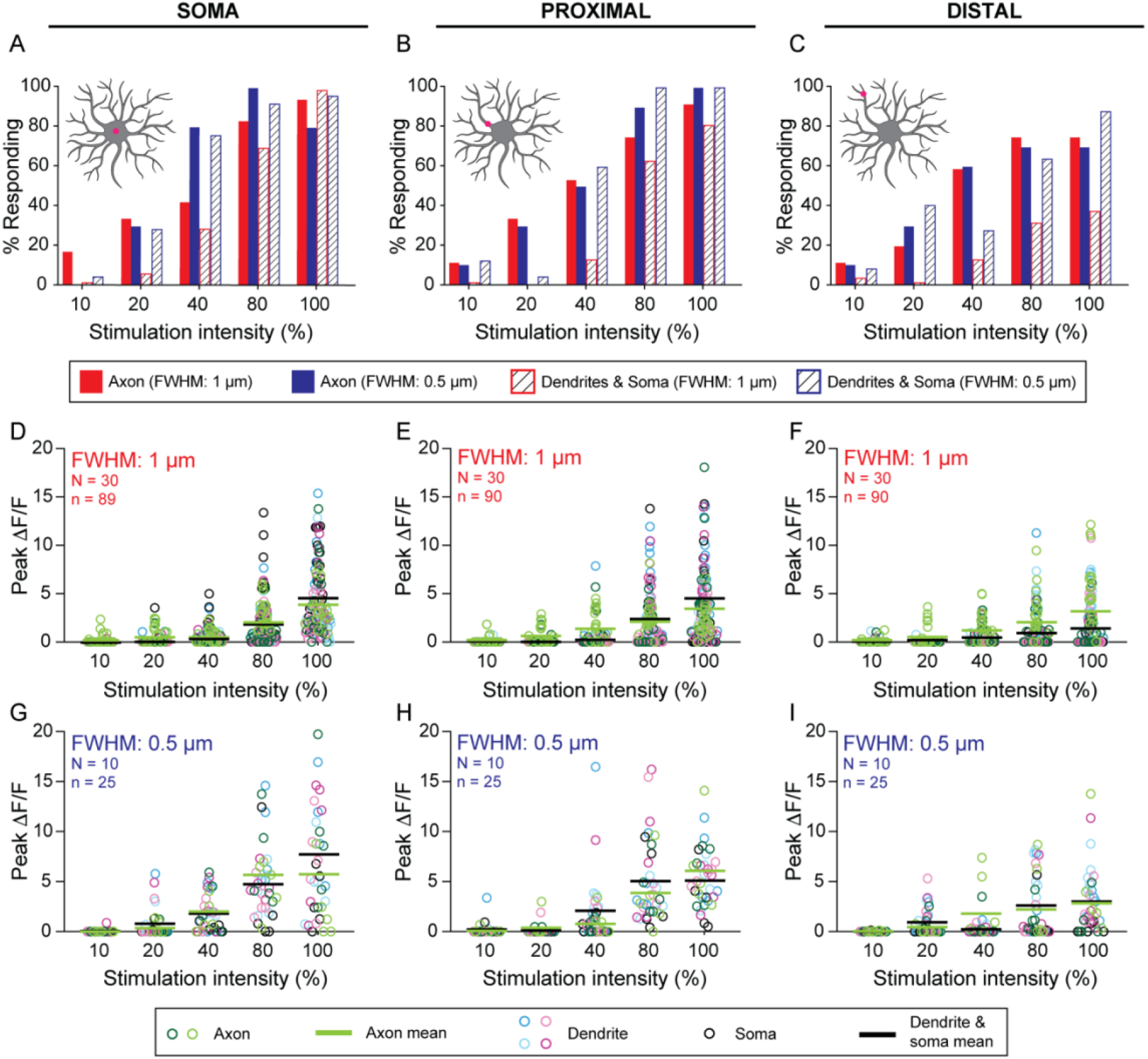
Characterization of the “contact” axonal and dendritic calcium response. (A-C) Fraction of cells responding to direct stimulation to the cell body (A), proximal dendrite (B), and distal dendrite (C). (D-I) Peak calcium responses in cells stimulated on the cell body (D,G), on proximal dendrites (E,H) and on distal dendrites (F,I). N represents number of larvae; n represents number of cells. Statistical analysis of these data are contained in *Supplementary Materials, Table 3-4*.

The magnitudes of the calcium responses (peak *ΔF*/*F*) increased with increasing stimulus intensity at all locations (Figure 4D-I, *Supplementary Figures 3-4, Supplementary Movie 5-6*). The average response magnitudes for somal and proximal stimulation were similar (Figure 4D,E,G,H) and somewhat larger than for distal stimulation (Figure 4F,I). The average axonal response magnitudes were similar to the average somal and dendritic magnitudes (green and black lines in Figures 4D-I).

Both the wider and narrower profiles triggered cellular calcium signals with similar frequency (Figure 4A-C). The magnitudes of the peak axonal responses were independent of the profile (Figure 4 D-I, green circles and bars), whereas the magnitudes of the dendritic and soma responses were significantly larger for the narrower profile (Figure 4 D-I, non-green circles and blacks bars). See *Supplementary Table 4* for statistical analysis. We will return to the question of how the stimulus and dendrite geometry effects the responses.

The latencies of the contact responses were shorter in the axons than the dendrites (Figure 5A,B): in other words, the dendritic response rises with a larger delay than the axonal response. For both axons and dendrites, higher intensities gave shorter latencies. The latencies of axonal contact responses were similar to those of non-contact responses (Figure 3E). Interestingly, the rising phases of the dendritic responses were almost simultaneous in all the dendritic regions, being within the 100 ms frame time of the camera, even though the latency was significantly longer (≥ 500 ms). For example, directly stimulating a peripheral dendrite gave a response in the same dendrite and in a dendrite on the other side of the cell body (>200 μm distance away) with a time-course that rose within 100 ms of each other (1 frame) (Figure 5C). This shows that the dendritic signals propagate at speeds >2 mm/s (= 200 μm /100 ms).

**Fig 5.**
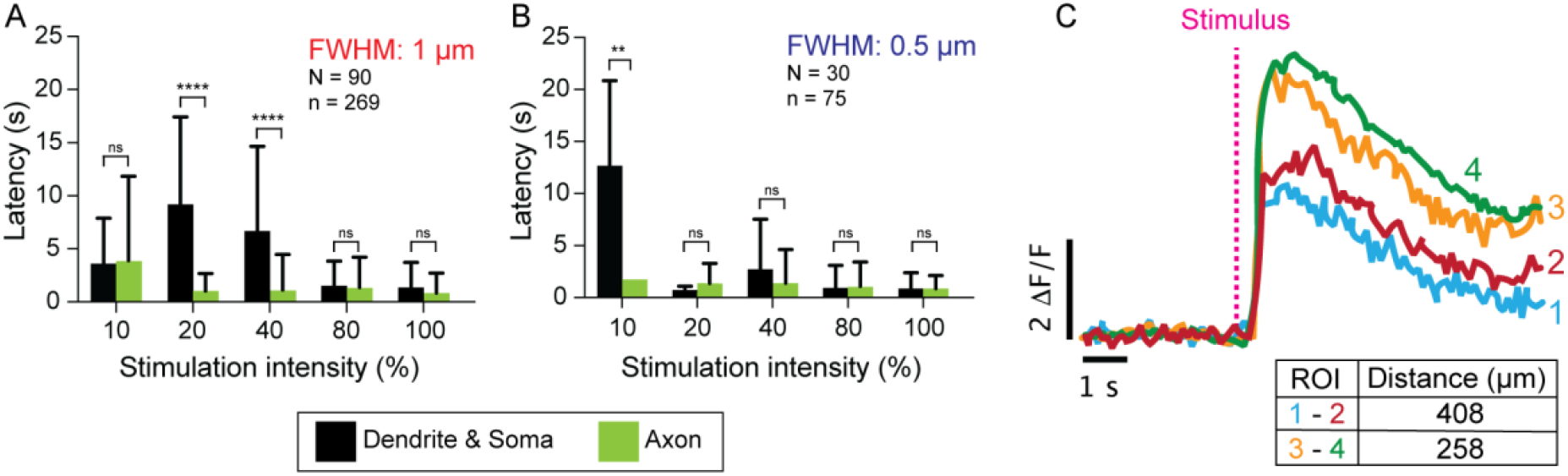
Latencies of axonal and dendritic responses to “contact” stimulation. (A-B) Latencies for cells stimulated with wider (A) and narrower (B) profiles at varying intensities (10-100%). For statistical comparisons across two different irradiance settings see *Supplementary Materials, Table 5*. (C) Representative traces from one neuron showing that dendritic regions >250 μm apart on opposite sides of the cell body respond simultaneously, though with a lag relative to the stimulus. See *Supplementary Materials Movie 4-6*.

### Contact responses depend on the overlap between the stimulus and the dendrite

The diameters of dendrites decrease from ∼1 μm for the proximal-most to ∼0.25 μm for the terminal dendrites (Liao and Howard, 2020). Therefore, both the narrower and wider stimuli will fall mostly within the proximal dendrites (and soma), whereas the wide stimulation will fall mostly outside the distal dendrites. Thus, wider profiles are expected to be less effective when applied distally. To test whether this accounts for the differences between proximal and distal stimulation (e.g. Figure 4F,I), we formulated a mathematical model that takes into consideration both the shape of the laser stimulus and the geometry of the dendrites. The laser beams were modelled as two-dimensional Gaussians with standard deviations corresponding to the measured FWHMs (Figure 6A). The dendrites were modelled as cylinders with the estimated diameters: 1 μm proximal and 0.4 μm distal (Figure 6B). The soma was modelled as a cylinder with diameter 2 μm. We accounted for possible non-linearity between stimulus intensity and response magnitude by introducing an exponentiation factor with exponent (*n*). We then asked whether the observed peak values of *ΔF*/*F* (replotted in Figure 6C-H) are consistent with the differing overlaps between the stimulus and the dendrite. In other words, does stimulation of thinner dendrites result in smaller responses because a smaller fraction of stimulus is actually making contact with the process? We considered two cases: (i) the response depends on the overlap of the stimulus with the dendrite volume and (ii) the response depends on the overlap of the stimulus with the dendrite surface area. The equations are in the *Materials and Methods*.

**Fig 6.**
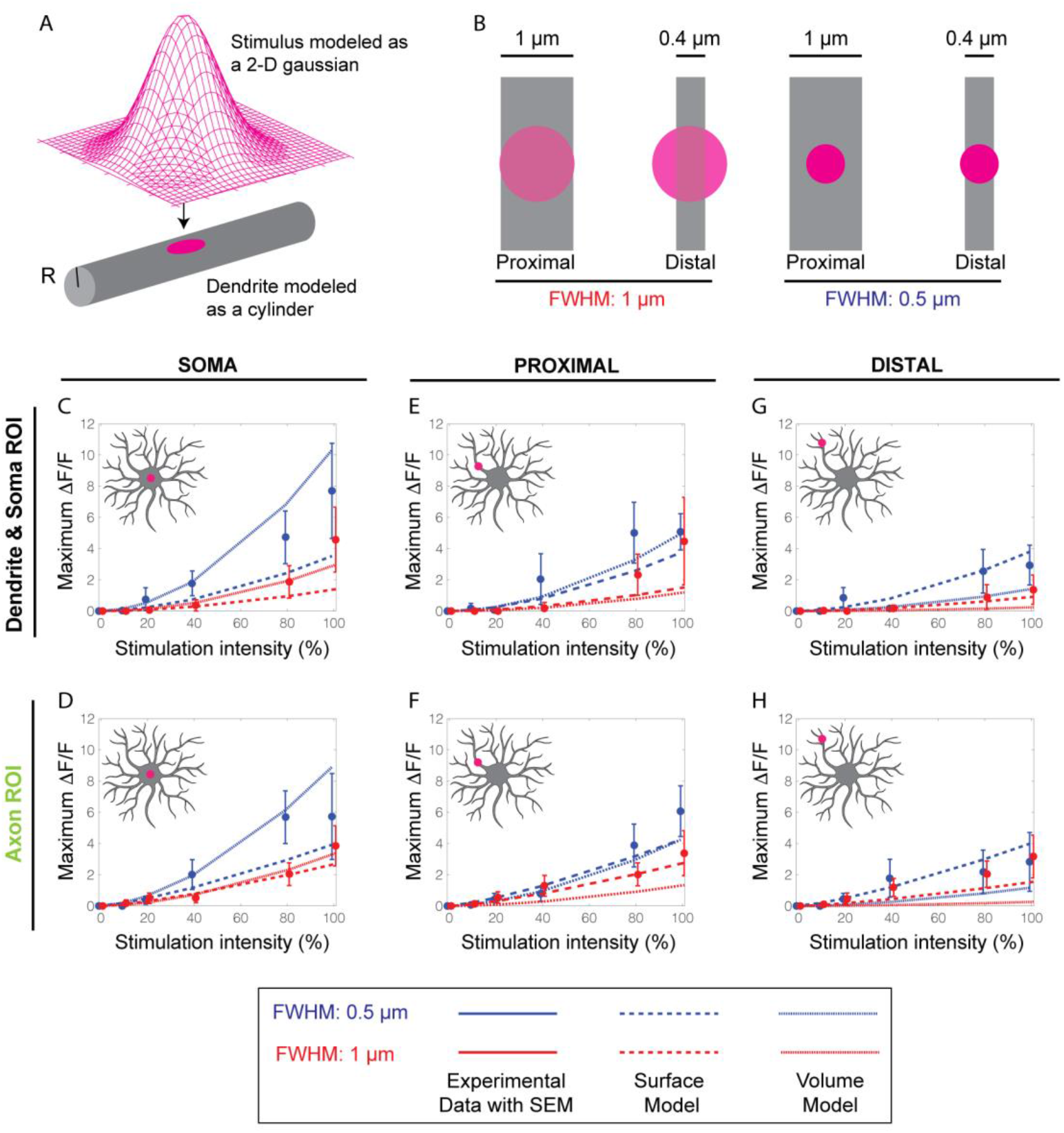
Overlap model for the effectiveness of stimuli in generating responses. (A) Schematic of the overlap model: the laser profile is approximated by a 2-D Gaussian and the dendrite modeled by a cylinder with radius *R*. (B) Top-down view of the two laser profiles projected onto proximal and distal dendrites. Proximal dendrites have radius 500 nm and distal dendrites have radius 200 nm. (C-H) Theoretical curves (lines) superimposed on the measured peak *ΔF*/*F* for somal (C, D), proximal dendrite (E, F) and distal dendrite (G, H) stimulation. Dashed lines represent the surface model and dotted line represents the volume model. Model parameters are listed in *Supplementary Table 6-7*.

We fit the models to the data (Figure 6C-H) using the measured profiles, dendrite diameters and stimulus intensities. For each model (volume, surface area) and data set (axon, dendrite/soma) we found the values the exponent (*n*) and a conversion factor (*γ*), that simultaneously minimized the sum of the least squares difference between model and all six associated experimental curves. The models recapitulated the non-linear increase in response with stimulus intensity, the observed smaller peak values of *ΔF*/*F* for distal stimulation, and the dependence on the profile widths. Interestingly, we found that the axonal responses were fit better to the surface model than the dendritic and somal responses. Thus, the responses depend on the overlap of the stimulus with the dendrite (*Supplementary Table 8*).

## Discussion

In this study we developed a non-lethal, tunable, *in vivo* assay for larval nociception using a 405-nm laser that causes highly localized puncture wounds to the larval cuticle. This stimulation evoked behavioral responses similar to nociceptive avoidance responses triggered by a wasp ovipositor. By tuning the intensity, duration, spatial profile and position of the laser focus, we could probe the conditions necessary for evoking calcium responses in Class IV neurons, which were monitored by increases in GCaMP6f fluorescence under a spinning disk confocal microscope.

Our primary finding is that there are two distinct calcium signaling responses in Class IV cells: (i) a non-contact response observed primarily in axons, and (ii) a contact response seen in axons, dendrites and cell bodies (Figure 7A). The existence of two response pathways is supported by three pieces of evidence: (a) axonal calcium signals do not require the laser spots to make direct contact with the dendritic processes (Figure 2A,B) whereas dendritic calcium signals require direct contact (Figure 2C,D); (b) axonal calcium signals are more sensitive and faster than dendritic calcium transients even when the stimulus is as far as 400 μm away from the axon (Fig 3, Fig 4); and (c) the surface model provides a better fit to the axon responses while the volume model provides a better fit to the dendrite and soma data (Fig 6C-H, *Supplementary Table 8*). Therefore, we tentatively conclude that localized mechanical damage induced by the laser triggers non-contact responses in the axons and contact responses in all cellular compartments. The conclusion that the axon-only response is indirect is strengthened by the observation that non-contact wide-profile illumination gives smaller axonal responses despite it having more power at larger distances (that could potentially directly stimulate the dendrite).

**Fig 7.**
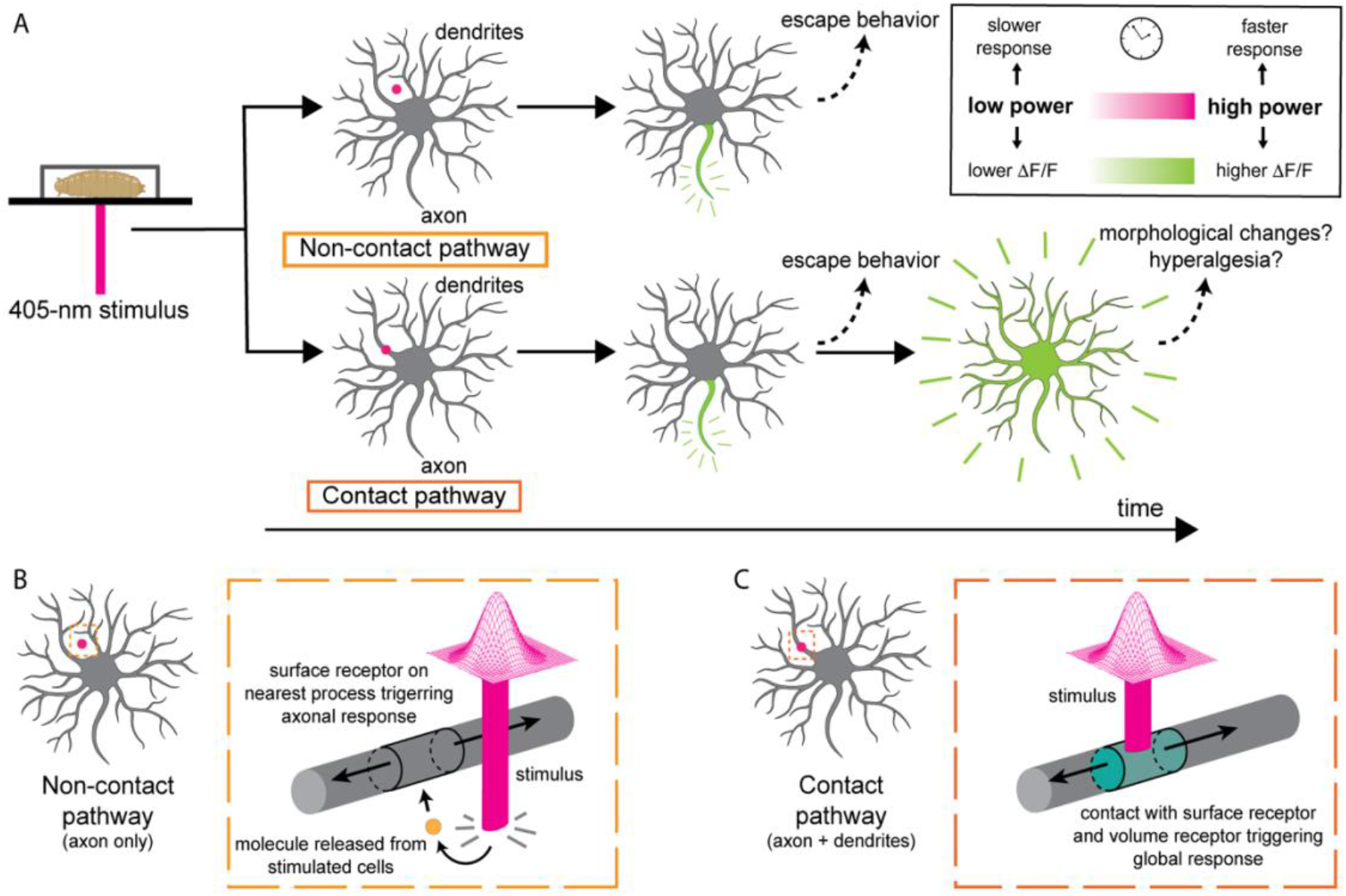
Summary of results and their interpretations. (A) Upper panels: non-contact stimulation (magenta dot) initiates axonal calcium responses. Lower panels: contact stimulation (magenta dot) initiates axonal and dendritic calcium responses. (B) Hypothetical mechanism underlying the non-contact response: damage to adjacent cells releases molecules (orange circle) that bind receptors on dendritic surface leading to cell depolarization. The depolarization is enough to trigger action potentials in the axon, which open calcium channels in the axon; the depolarization is insufficient to open calcium channels in the dendrites and soma. (C) Hypothetical mechanism underlying the contact response: direct damage to the dendrite strongly depolarizes the cell and opens calcium channels in dendrites, soma and axon. The contact stimulus is also expected to also trigger the non-contact pathway.

Given that stimulation with a focused laser shares several features with stimulation by an ovipositor – localized tissue damage, melanotic spots, behavioral responses, axonal signals – we postulate that the ovipositor can excites the Class IV neuron through both the contact and non-contact mechanisms. There are, however, some potential caveats to this conclusion. First, a wasp ovipositor punctures the cuticle via mechanical pressure, whereas our laser is likely damaging the cuticle via localized heating or production of reactive oxygen species by autofluorescence or GCaMP6f fluorescence. Second, while both the ovipositor and the focused laser produce localized damage, they are both expected to produce more delocalized effects on the tissue. The ovipositor is expected to generate a large strain field as the cuticle is indented before it ruptures. This strain field could evoke mechanoreceptive responses. The laser generates stray light over a wide area of the tissue through reflection and scattering, though the intensity is greatly attenuated. This stray light could excite photoreceptors (Xiang et al., 2010) or the reactive-oxygen-species response (Kim and Johnson, 2014). However, the stray light evidently does not excite dendritic calcium responses. Despite differences between laser and ovipositor stimulation, and the considerable uncertainty about the precise effects of ovipositor penetration and laser illumination on the tissue, we believe that the ovipositor likely stimulates both contact and non-contact responses.

We propose the following pathways to account for the contact and non-contact calcium responses. First, we propose that direct contact of high-power laser illumination damages the Class IV cell’s plasma membrane, making it more permeable to sodium and inducing a local depolarization of the membrane potential (Tracey, 2017). The depolarization then spreads electrotonically throughout the dendrite, to the cell body and the axon. Modeling electrotonic spread in the thin axons of primate rods and cones (which have diameters 0.45 and 1.6 μm respectively), shows that there is little signal decrement over 400 μm even at frequencies up to 50 Hz, which corresponds to a time constant <10 ms (Hsu et al., 1998). Therefore, electrotonic spread of depolarization is likely fast enough to reach all parts of the Class IV cell. If the depolarization exceeds the threshold needed to open L-type (and potentially other) calcium channels, then calcium will enter and a GCaMP6f fluorescent signal produced. If there are calcium channels in the dendrites, cell body and axon, then fluorescence changes will be observed throughout the cell.

Second, we propose that if high-power laser illumination makes no contact with the Class IV cell, it will, never-the-less damage adjacent cells, such as the overlying epithelial (epidermal) cells and underlying muscle cells (Grueber et al., 2002). These cells could then release small metabolites or acidify the extracellular space. These signals then spread by diffusion to the membrane of the Class IV cells where they open receptor-gated or the acid-sensing channels — for example, pickpocket or ripped pocket (Adams et al., 1998; Boiko et al., 2011). This mechanism would be analogous to release of cytosolic ATP from damaged cells, which mediates pain perception via contact with P2X receptors on peripheral nociceptive cells in vertebrates (Cook and McCleskey, 2002; Hamilton and McMahon, 2000). While *Drosophila* lacks P2X receptors (Fountain and Burnstock, 2009), it is possible that other small cytoplasmic molecules or protons released by surrounding cells might play an analogous role. Opening of receptor-linked channels is expected to locally depolarize the cell membrane, and this depolarization will spread electrotonically to the cell body and axon, where, if it exceeds a threshold, leads to axonal action potentials which in turn trigger the opening of calcium channels. If the receptor mechanism leads to less depolarization in Class IV dendrites than direct damage, as is reasonable, then non-contact stimulation may be above threshold for action potentials in the axons (which then open calcium channels), but below threshold for opening calcium channels in the dendrites and soma. Hence, only axons respond to non-contact stimulation. Because direct contact is also likely to damage adjacent cells and trigger the non-contact response as well, axon responses are likely to be triggered by both pathways. Thus, there are likely two pathways by which localized damage by ovipositor barbs leads to electrophysiological and calcium responses.

Interestingly, the existence of these two pathways provides evidence that the dendrites of Class IV cell are not electrically excitable. If they were excitable, then we would expect that axonal action potentials would back propagate and in turn stimulate calcium entry through voltage-gated channels in the dendrites; but the non-contact response does not stimulate calcium responses in dendrites. A related point is that when direct contact is made, the axonal calcium signals (*Supplementary Figure* 4) are usually more transient than the dendritic signals (*Supplementary Figure* 3). A possible explanation is that calcium entry opens calcium-activated potassium channels in the axons, which depolarizes the axonal membrane tending to inhibit spiking and additional calcium entry. This delayed negative feedback would attenuate the calcium signal in the axon at longer times. The existence of axonal calcium-activated potassium channels could account for the “unconventional spikes” (US) recorded from the cell body and the axon bundle (Terada et al., 2016): these spikes are characterized by an ensuing refractory period during which there is no spiking; the US and refractory period correlates with calcium signals in the dendrites and may be a consequence of the opening of calcium-activated potassium channels.

The existence of the non-contact pathway sheds new light on the highly branched morphology of Class IV cells. Because the “mesh size” — the average distance between dendrites in the arbor — is about 5 μm, it has been suggested that the reason these cells are highly branched is to maximize direct contact with ovipositor barbs (Ganguly et al., 2016). However, the non-contact pathway implies that direct damage to the Class IV cell is not necessary to stimulate the axonal pathway. However, the Class IV cells still need to be highly branched and make a fine mesh so that extracellular signals can still diffuse sufficiently quickly to activate membrane receptors: a small molecule similar in size to ATP (diffusion coefficient on the order of 100 μm ^2^/sec) will reach a dendrite 5 μm away in ∼0.1 second. To diffuse a distance three times as far (15 μm) would take ∼1 second, too slow to account for the axonal responses. Thus, our data lead us to propose a new function underlying extensive branching of Class IV dendritic arbors: the fine meshwork minimizes diffusion times to ensure that non-contact stimulation is rapidly transduced.

While the function of the axonal response is clear – to convey nociceptive signals to the central nervous system, the function of the dendritic is not. The dendritic signals are often centrifugal, moving away from the cell body; they are therefore not on the cell-to-brain pathway. One implication is that dendritic calcium signals in Class IV cells are not necessarily good proxies for neuronal excitation. Calcium signals are often assumed to be reporters of cell excitation, though a number of researchers have cautioned against this assumption (Ali and Kwan, 2019; Higley and Sabatini, 2008). It is possible that dendritic calcium mediates hyperalgesia by modifying the sensitivity in case of a second attack. Alternatively, because severing dendrites leads to peripheral degeneration (Song et al., 2012), it is possible that the dendrite-wide calcium signal could promote regrowth. These will be important possibilities to follow up on in future experiments.

## Materials and Methods

### *Drosophila* Strains and Husbandry

Fly lines were obtained from the Bloomington *Drosophila* Stock Center and through generous gifts from Damon Clark and Fernando Von Hoff. Fly stocks were maintained at 25°C in a humidity-controlled incubator (60% humidity) on standard apple-agar based food (Archon Scientific) with 12 -hour light/dark cycles. Fly crosses were maintained in fly chambers on apple juice agar-based food (mixture of apple agar concentrate, propionic acid, phosphoric acid and water) with a generous dollop of yeast paste at 25°C, 60% humidity. Larvae 68-72 hours after egg laying were used for all imaging experiments. The following fly lines were used to image Class IV da neurons:

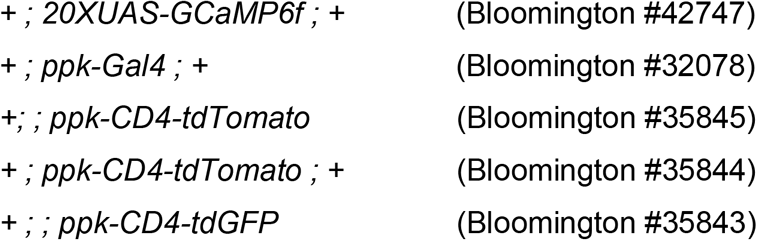

### Microscopy and imaging experiments

#### Live-cell Confocal Imaging

Larvae were timed and selected 68-72 hours AEL for imaging. Prior to imaging, larvae were washed in distilled water and gently rolled on a glass slide with a paintbrush to remove excess food and debris. Larvae were then placed on a Cellvis 35mm glass bottom dish (D35-20-1.5-N) and allowed to acclimatize for 60 seconds. Larvae were then immobilized using a single-layer PDMS device using a protocol as previously described (Mishra et al., 2014). Briefly, larvae were positioned on the center of the dish and gently constrained inside the PDMS cavity. The PDMS was then adhered to the dish by applying slight suction using a 30ml syringe. No anesthetic was used. Samples were then mounted on the microscope stage, illuminated with Nikon lasers (488nm or 561nm at 30-50% laser power) and imaged at 8-10 Hz on a spinning disk microscope: Yokogawa CSU-W1 disk (pinhole size 50 μm) built on a fully automated Nikon TI inverted microscope with perfect focus system, an sCMOS camera (Zyla 4.2 plus sCMOS), and Nikon Elements software with either a 40X (1.25 NA, 0.1615 micron pixel size) or 20X (0.50 NA, 0.3225 micron pixel size). The temperature of the sample region was maintained using an objective space heater at 25°C. Samples were manually focused for each cell prior to image acquisition. No more than 3 cells were imaged from an individual larval sample. All data sets represent cells from at least four independent larval samples.

#### 405-nm Stimulation

Stimulation of Class IV da neurons was performed using a 405-nm laser (OBIS 405 nm LX 100 mW, Coherent, Santa Clara, CA) which was connected to the microscope through an empty port. Integrated wattage values of the laser were measured using a microscope slide power sensor (Thor Labs, Newton, NJ) at the sample plane. Activation of the laser was synchronized to the imaging rate using a custom LabView macro. Stimulus intensity was user-defined before each experiment (10-100%, 0-43 mW integrated power) and administered for 100ms. The precise location of the laser was calibrated using a custom graticule set in NIS Elements (Nikon) and tested prior to each experiment. For images targeting the soma, the laser was focused on the center of the cell body. Proximal dendrites were stimulated along a main branch 10-30 μm from the cell body. For distal branches, stimulus was administered to a branch 150-200 μm from the soma. Stimulation experiments were performed over 30-45 seconds wherein the stimulus was administered after 10-12 seconds of initial baseline recording for each cell.

### Data Analysis

#### Image Processing

Movies were analyzed using Image J (NIH). When necessary, movies were stabilized using the Template-Matching or Image Stabilizer plug-ins. For each cell, several regions of interest (ROI) were manually selected for each cell from 7 different locations along the entire dendritic tree to study any differential responses within the same cell: soma (1 ROI), axon (2 ROIs), dendritic arbors (4 ROIs). Care was taken minimize background by contouring the ROI region to encompass only the cellular region being considered. Corresponding fluorescence values for each ROI were extracted in Image J and imported into MATLAB (Mathworks). Baseline fluorescence *F*_0_ was calculated as the camera’s mean fluorescence signal for all frames before laser stimulation. The difference in fluorescence from the baseline, (*ΔF*/*F*), was calculated as (*F* − *F*_0_)/(*F*_0_ − 100) where is the fluorescence signal and 100 is the manufacturer’s camera offset. The time-series data were median filtered (width 7) to remove outliers resulting from noise or movement. For measurement of puncture wounds, cells were stimulated and then z-stacks were acquired at 0.5 μm z-intervals. Diameters were then analyzed by taking line scans through the center of the wounds on maximum-projection images.

#### Calcium Imaging Response Criteria

ROIs were scored as being responsive to the stimulus if the *ΔF*/*F* at any frame after stimulation was greater than 5 standard deviations above the baseline before stimulation. The largest *ΔF*/*F* value for all frames post stimulation was determined to be peak values *ΔF*/*F*. The timepoint when *ΔF*/*F* was equal to or greater than 5 standard deviations above baseline *F* was defined as the latency.

#### Modeling

We modeled the observed dendritic calcium signal magnitudes as a function of intensity (integrated power) and irradiance. Can the overlapping geometry of the stimulus and the dendrite account for the observed differences in responses to narrow and wide stimulus profiles and when applied to thick, proximal dendrites and thin, distal dendrites? The laser was modeled as a two-dimensional Gaussian with experimentally measured variance: the spatial profile of the laser was experimentally measured using interference-reflection microscopy (MAHAMDEH et al., 2018) by analyzing the reflection of the laser on a coverslip and using a line scan in ImageJ. A Gaussian was fit over the line scan in MATLAB to compute the standard deviation σ. To test whether the observed laser-activated calcium responses are due to surface or volume illumination, we considered two different models. First, we considered overlap of the laser profile with the cylindrical surface of the dendrite (Eq 1). In the second model, we considered overlap of the laser profile with the cylindrical volume of the dendrite (Eq 2).

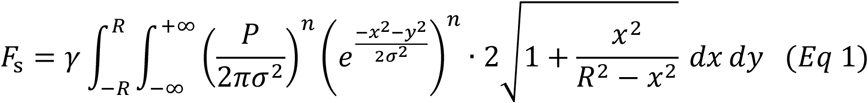

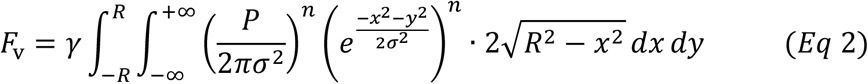

Here, *F*_s_ and *F*_v_ are the theoretical peak values of *ΔF*/*F* corresponding to each model, *P* is the integrated laser power, σ is the standard deviation of the Gaussian, and *R* is the radius of the dendrite. The soma was modeled as a cylinder with radius *R*_soma_ = 1 μm significantly larger than that of proximal (0.5 μm) and distal dendrites (0.2 μm) (Liao and Howard, 2020). This simplification is justified as the laser profile dies off exponentially and σ ≪ *R*_soma_. The variable *n* is a free parameter introduced to account for the observed nonlinearity in experimental values, and *γ* is a free parameter corresponding to a conversion factor between units.

*P* ranged between 0 to 100 based on the power output of the laser. σ was set at 212.31 nm or 424.62 nm, corresponding to the two different stimulation irradiance settings. Because peak values of *ΔF*/*F* exhibited unequal variances (heteroskedasticity) across the range of stimulation wattages, we computed a set of weights for use in our weighted least squares fitting by performing a linear regression between the *ΔF*/*F* and the experimental SEM. A detailed table of input values for the models can be found in the Supplementary Materials. MATLAB’s *fminsearch* was used to compute the values for *n* and *γ* that simultaneously minimized the sum of the squared errors between all theoretical and experimental values. Minimization was performed by considering data from axon ROIs and non-axon ROIs separately. The surface and volume model were each fit to the data.

### Statistical Analysis

Sample sizes are listed for each data set on the corresponding plots. Capitalized ‘N’ indicates number of larvae; lowercase n is number of neurons. Statistical analysis was performed in Prism 8 (GraphPad). Sidak’s test was used when making pairwise comparisons of multiple comparisons. One-way analysis of variance (ANOVA) was used to determine if statistically significant differences existed between the means of three or more independent groups. For plots showing peak values of *ΔF*/*F*, all data points (open circles) and experimental means (lines) are shown on graphs to demonstrate experimental variability. For plots showing latency, experimental means and SD are shown. Significance was evaluated at P < 0.05.

## Acknowledgements

We thank all members of the Howard lab for many helpful discussions and encouragement. Special thanks to Drs. Mohammed Mahamdeh for building the laser set up on the confocal microscope, and to Sonal Shree and Anna Luchniak for technical assistance. The authors also acknowledge Drs. Damon Clark, Yong Xiong, Fernando Von Hoff, Sean Christie (MVI) for technical and scientific advice. Some images in this manuscript were modified from BioRender.

## Author Contributions

R.B and J.H conceptualized all experiments and models. R.B performed all experiments.

R.B and S.S performed all analysis. R.B, S.S, and J.H prepared the manuscript.

## Point of Contact

Any inquiries should be directed to Jonathon Howard.

## Supplementary Materials

**Table 1:** Comparison of magnitude of responses shown in Figure 3 C, D. Sidak’s multiple comparisons test results shown for comparisons.

|  | Figure 3C | Figure 3C | Figure 3C, D | Figure 3C, D |
| --- | --- | --- | --- | --- |
| | Axon (green line)<br>(FWHM: 1 $\mu\text{m}$)<br>vs.<br>Dendrite & Soma<br>(black line)<br>(FWHM: 1 $\mu\text{m}$) | Axon (green line)<br>(FWHM: 0.5 $\mu\text{m}$)<br>vs.<br>Dendrite & Soma<br>(black line)<br>(FWHM: 0.5 $\mu\text{m}$) | Axon (green line)<br>(FWHM: 1 $\mu\text{m}$)<br>vs.<br>Axon (green line)<br>(FWHM: 0.5 $\mu\text{m}$) | Dendrite & Soma<br>(black line)<br>(FWHM: 1 $\mu\text{m}$)<br>vs.<br>Dendrite & Soma<br>(black line)<br>(FWHM: 0.5 $\mu\text{m}$) |
| 10% | p = 0.9317<br>(ns) | p = 0.1117<br>(ns) | p = 0.9751<br>(ns) | p = 0.8456<br>(ns) |
| 20% | P = 0.7630<br>(ns) | p = 0.0175<br>(*) | p = 0.7790<br>(ns) | p = 0.9719<br>(ns) |
| 40% | p < 0.0001<br>(****) | p < 0.0001<br>(****) | p = 0.9896<br>(ns) | p = 0.1417<br>(ns) |
| 80% | p < 0.0001<br>(****) | p < 0.0001<br>(****) | p > 0.9999<br>(ns) | p = 0.0032<br>(**) |
| 100% | p < 0.0001<br>(****) | p < 0.0001<br>(****) | p > 0.9999<br>(ns) | p = 0.0193<br>(**) |

**Table 2:** Comparison of latencies for axon ROIs stimulated under the “non-contact” condition as shown in Figure 3E. Sidak’s multiple comparisons test results shown.

|  | **Figure 3E** |
| --- | --- |
|  | **Axonal**<br>**Response**<br>**latency**<br>**(FWHM: 1 μm)**<br>**vs.**<br>**(FWHM: 1 μm)** |
| **10%** | p = 0.2011<br>(ns) |
| **20%** | p = 0.0004<br>(***) |
| **40%** | p = 0.4851<br>(ns) |
| **80%** | p = 0.9710<br>(ns) |
| **100%** | p > 0.9999<br>(ns) |

**Table 3:** Comparison of magnitude of responses between axons and dendrite/soma ROIs for the “contact” dendritic response as shown in Figures 4 D-I. Sidak’s multiple comparisons test results shown for comparisons.

|  | **Figure 4D**<br>Soma Stim | **Figure 4E**<br>Prox Stim | **Figure 4F**<br>Distal Stim | **Figure 4G**<br>Soma Stim | **Figure 4H**<br>Prox Stim | **Figure 4I**<br>Distal Stim |
| --- | --- | --- | --- | --- | --- | --- |
| | **Axon (green line)**<br>(FWHM: 1 μm)<br>vs.<br>**Dendrite & Soma (black line)**<br>(FWHM: 1 μm) | **Axon (green line)**<br>(FWHM: 1 μm)<br>vs.<br>**Dendrite & Soma (black line)**<br>(FWHM: 1 μm) | **Axon (green line)**<br>(FWHM: 1 μm)<br>vs.<br>**Dendrite & Soma (black line)**<br>(FWHM: 1 μm) | **Axon (green line)**<br>(FWHM: 0.5 μm)<br>vs.<br>**Dendrite & Soma (black line)**<br>(FWHM: 0.5 μm) | **Axon (green line)**<br>(FWHM: 0.5 μm)<br>vs.<br>**Dendrite & Soma (black line)**<br>(FWHM: 0.5 μm) | **Axon (green line)**<br>(FWHM: 0.5 μm)<br>vs.<br>**Dendrite & Soma (black line)**<br>(FWHM: 0.5 μm) |
| **10%** | p = 0.9975<br>(ns) | p = 0.9998<br>(ns) | p = 0.9998<br>(ns) | p > 0.9999<br>(ns) | p > 0.9999<br>(ns) | p > 0.9999<br>(ns) |
| **20%** | p = 0.8916<br>(ns) | p = 0.9006<br>(ns) | p = 0.6027<br>(ns) | p = 0.9993<br>(ns) | p = 0.9991<br>(ns) | p = 0.9904<br>(ns) |
| **40%** | p = 0.9993<br>(ns) | p = 0.2950<br>(ns) | p = 0.0041<br>(**) | p > 0.9999<br>(ns) | p = 0.6824<br>(ns) | p = 0.2377<br>(ns) |
| **80%** | p = 0.9981<br>(ns) | p = 0.9921<br>(ns) | p = 0.0010<br>(***) | p = 0.9593<br>(ns) | p = 0.7610<br>(ns) | p = 0.9954<br>(ns) |
| **100%** | p = 0.4851<br>(ns) | p = 0.3031<br>(ns) | p = <0.0001<br>(****) | p = 0.5376<br>(ns) | p = 0.8493<br>(ns) | p > 0.9999<br>(ns) |

**Table 4:** Comparison of magnitude of responses shown in Figure 4 D-I across two different stimulation irradiance settings. Axon ROI (FWHM: 1 μm) were compared to axon ROI (FWHM: 0.5 μm). Dendrite & soma ROI (FWHM: 1 μm) were compared to dendrite & soma ROI (FWHM: 0.5 μm). Sidak’s multiple comparisons test results shown for comparisons.

|  | Figure 4D,G<br>Soma Stim | Figure 4D,G<br>Prox Stim | Figure 4E,H<br>Distal Stim | Figure 4E,H<br>Soma Stim | Figure 4F,I<br>Prox Stim | Figure 4F,I<br>Distal Stim |
| --- | --- | --- | --- | --- | --- | --- |
| | Axon (green line) (FWHM: 1 μm) vs. Axon (green line) (FWHM: 0.5 μm) | Axon (green line) (FWHM: 1 μm) vs. Axon (green line) (FWHM: 0.5 μm) | Axon (green line) (FWHM: 1 μm) vs. Axon (green line) (FWHM: 0.5 μm) | Dendrite & Soma (black line) (FWHM: 1 μm) vs. Dendrite & Soma (black line) (FWHM: 0.5 μm) | Dendrite & Soma (black line) (FWHM: 1 μm) vs. Dendrite & Soma (black line) (FWHM: 0.5 μm) | Dendrite & Soma (black line) (FWHM: 1 μm) vs. Dendrite & Soma (black line) (FWHM: 0.5 μm) |
| 10% | p = 0.9998<br>(ns) | p > 0.9999<br>(ns) | p > 0.9999<br>(ns) | p > 0.9999<br>(ns) | p = 0.9998<br>(ns) | p > 0.9999<br>(ns) |
| 20% | p > 0.9999<br>(ns) | p = 0.9999<br>(ns) | p > 0.9999<br>(ns) | p = 0.8078<br>(ns) | p > 0.9999<br>(ns) | p = 0.0675<br>(ns) |
| 40% | p = 0.2251<br>(ns) | p = 0.9626<br>(ns) | p = 0.9434<br>(ns) | p = 0.1265<br>(ns) | p = 0.0605<br>(ns) | p > 0.9999<br>(ns) |
| 80% | p < 0.0001<br>(****) | p < 0.0563<br>(ns) | p > 0.9999<br>(ns) | p < 0.0001<br>(****) | p = 0.0016<br>(**) | p < 0.0001<br>(****) |
| 100% | p = 0.0746<br>(ns) | p = 0.0012<br>(**) | p = 0.9940<br>(ns) | p < 0.0001<br>(****) | p = 0.9380<br>(ns) | p < 0.0001<br>(****) |

**Table 5:** Comparison of latencies for ROIs stimulated with two different irradiance settings in Figure 5. Sidak’s multiple comparisons test results shown.

|  | Figure 5A, B | Figure 5A, B |
| --- | --- | --- |
| | **Dendrite & Soma (black)**<br>(FWHM: 1 $\mu\text{m}$)<br>vs.<br>**Dendrite & Soma (black)**<br>(FWHM: 0.5 $\mu\text{m}$) | **Axon (green)**<br>(FWHM: 1 $\mu\text{m}$)<br>vs.<br>**Axon (green)**<br>(FWHM: 0.5 $\mu\text{m}$) |
| **10%** | $p = 0.0052$<br>(**) | $p = 0.9994$<br>(ns) |
| **20%** | $p < 0.0001$<br>(****) | $p > 0.9999$<br>(ns) |
| **40%** | $p < 0.0001$<br>(****) | $p > 0.9999$<br>(ns) |
| **80%** | $p = 0.9046$<br>(ns) | $p > 0.9999$<br>(ns) |
| **100%** | $p = 0.9537$<br>(ns) | $p > 0.9999$<br>(ns) |

**Table 6:**
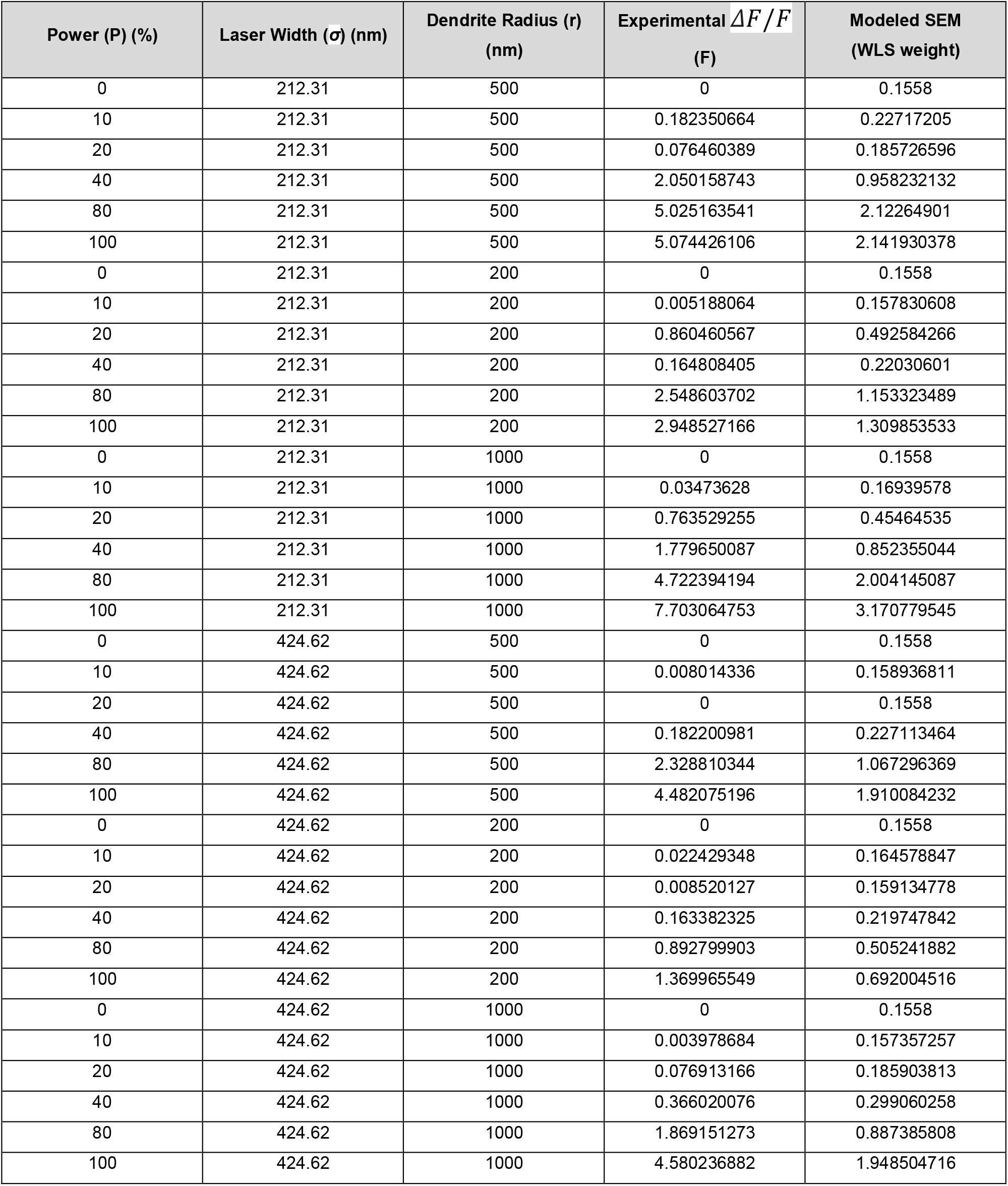
Modeling input parameter values for dendrite and soma ROI. 100% power corresponds to 43 mW for both laser profiles (see Supplementary Figure 1).

| Power (P) (%) | Laser Width ($\sigma$) (nm) | Dendrite Radius (r) (nm) | Experimental $\Delta F/F$ (F) | Modeled SEM (WLS weight) |
| --- | --- | --- | --- | --- |
| 0 | 212.31 | 500 | 0 | 0.1558 |
| 10 | 212.31 | 500 | 0.182350664 | 0.22717205 |
| 20 | 212.31 | 500 | 0.076460389 | 0.185726596 |
| 40 | 212.31 | 500 | 2.050158743 | 0.958232132 |
| 80 | 212.31 | 500 | 5.025163541 | 2.12264901 |
| 100 | 212.31 | 500 | 5.074426106 | 2.141930378 |
| 0 | 212.31 | 200 | 0 | 0.1558 |
| 10 | 212.31 | 200 | 0.005188064 | 0.157830608 |
| 20 | 212.31 | 200 | 0.860460567 | 0.492584266 |
| 40 | 212.31 | 200 | 0.164808405 | 0.22030601 |
| 80 | 212.31 | 200 | 2.548603702 | 1.153323489 |
| 100 | 212.31 | 200 | 2.948527166 | 1.309853533 |
| 0 | 212.31 | 1000 | 0 | 0.1558 |
| 10 | 212.31 | 1000 | 0.03473628 | 0.16939578 |
| 20 | 212.31 | 1000 | 0.763529255 | 0.45464535 |
| 40 | 212.31 | 1000 | 1.779650087 | 0.852355044 |
| 80 | 212.31 | 1000 | 4.722394194 | 2.004145087 |
| 100 | 212.31 | 1000 | 7.703064753 | 3.170779545 |
| 0 | 424.62 | 500 | 0 | 0.1558 |
| 10 | 424.62 | 500 | 0.008014336 | 0.158936811 |
| 20 | 424.62 | 500 | 0 | 0.1558 |
| 40 | 424.62 | 500 | 0.182200981 | 0.227113464 |
| 80 | 424.62 | 500 | 2.328810344 | 1.067296369 |
| 100 | 424.62 | 500 | 4.482075196 | 1.910084232 |
| 0 | 424.62 | 200 | 0 | 0.1558 |
| 10 | 424.62 | 200 | 0.022429348 | 0.164578847 |
| 20 | 424.62 | 200 | 0.008520127 | 0.159134778 |
| 40 | 424.62 | 200 | 0.163382325 | 0.219747842 |
| 80 | 424.62 | 200 | 0.892799903 | 0.505241882 |
| 100 | 424.62 | 200 | 1.369965549 | 0.692004516 |
| 0 | 424.62 | 1000 | 0 | 0.1558 |
| 10 | 424.62 | 1000 | 0.003978684 | 0.157357257 |
| 20 | 424.62 | 1000 | 0.076913166 | 0.185903813 |
| 40 | 424.62 | 1000 | 0.366020076 | 0.299060258 |
| 80 | 424.62 | 1000 | 1.869151273 | 0.887385808 |
| 100 | 424.62 | 1000 | 4.580236882 | 1.948504716 |

**Table 7:**
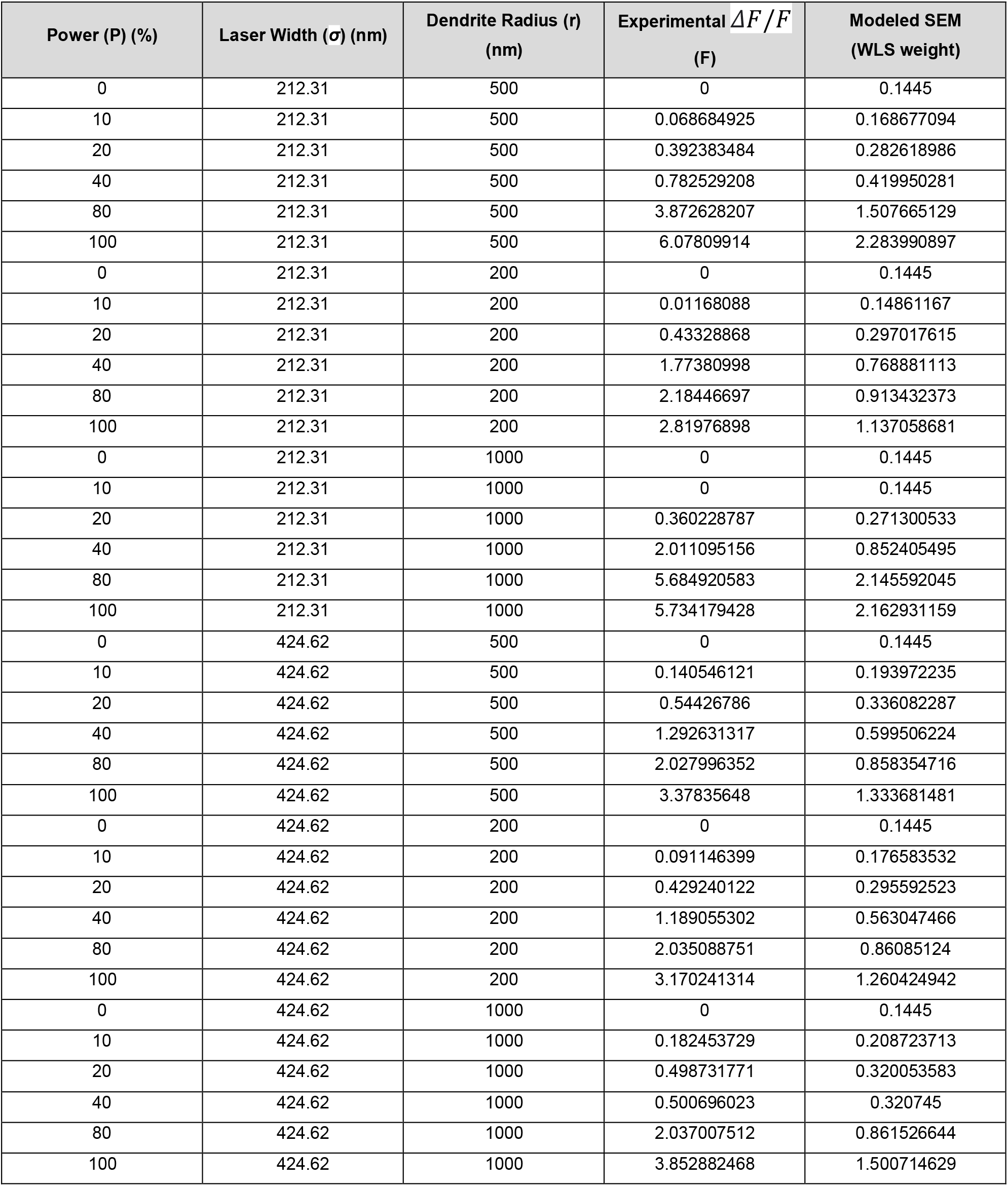
Modeling input parameter values for axon ROIs. 100% power corresponds to 43 mW for both laser profiles (see Supplementary Figure 1).

| Power (P) (%) | Laser Width ($\sigma$) (nm) | Dendrite Radius (r) (nm) | Experimental $\Delta F/F$ (F) | Modeled SEM (WLS weight) |
| --- | --- | --- | --- | --- |
| 0 | 212.31 | 500 | 0 | 0.1445 |
| 10 | 212.31 | 500 | 0.068684925 | 0.168677094 |
| 20 | 212.31 | 500 | 0.392383484 | 0.282618986 |
| 40 | 212.31 | 500 | 0.782529208 | 0.419950281 |
| 80 | 212.31 | 500 | 3.872628207 | 1.507665129 |
| 100 | 212.31 | 500 | 6.07809914 | 2.283990897 |
| 0 | 212.31 | 200 | 0 | 0.1445 |
| 10 | 212.31 | 200 | 0.01168088 | 0.14861167 |
| 20 | 212.31 | 200 | 0.43328868 | 0.297017615 |
| 40 | 212.31 | 200 | 1.77380998 | 0.768881113 |
| 80 | 212.31 | 200 | 2.18446697 | 0.913432373 |
| 100 | 212.31 | 200 | 2.81976898 | 1.137058681 |
| 0 | 212.31 | 1000 | 0 | 0.1445 |
| 10 | 212.31 | 1000 | 0 | 0.1445 |
| 20 | 212.31 | 1000 | 0.360228787 | 0.271300533 |
| 40 | 212.31 | 1000 | 2.011095156 | 0.852405495 |
| 80 | 212.31 | 1000 | 5.684920583 | 2.145592045 |
| 100 | 212.31 | 1000 | 5.734179428 | 2.162931159 |
| 0 | 424.62 | 500 | 0 | 0.1445 |
| 10 | 424.62 | 500 | 0.140546121 | 0.193972235 |
| 20 | 424.62 | 500 | 0.54426786 | 0.336082287 |
| 40 | 424.62 | 500 | 1.292631317 | 0.599506224 |
| 80 | 424.62 | 500 | 2.027996352 | 0.858354716 |
| 100 | 424.62 | 500 | 3.37835648 | 1.333681481 |
| 0 | 424.62 | 200 | 0 | 0.1445 |
| 10 | 424.62 | 200 | 0.091146399 | 0.176583532 |
| 20 | 424.62 | 200 | 0.429240122 | 0.295592523 |
| 40 | 424.62 | 200 | 1.189055302 | 0.563047466 |
| 80 | 424.62 | 200 | 2.035088751 | 0.86085124 |
| 100 | 424.62 | 200 | 3.170241314 | 1.260424942 |
| 0 | 424.62 | 1000 | 0 | 0.1445 |
| 10 | 424.62 | 1000 | 0.182453729 | 0.208723713 |
| 20 | 424.62 | 1000 | 0.498731771 | 0.320053583 |
| 40 | 424.62 | 1000 | 0.500696023 | 0.320745 |
| 80 | 424.62 | 1000 | 2.037007512 | 0.861526644 |
| 100 | 424.62 | 1000 | 3.852882468 | 1.500714629 |

**Table 8:** Summary of modeling fit parameters for dendrite and soma ROI and axon ROIs using the surface model (equation 1, *See Materials and Methods*) and volume model (equation 2, *See Materials and Methods*). Input parameters for the model are provided separately in Table 6 and Table 7. Visual plots of modeling results are shown in Figure 6 C-H.

|  | Surface Model (Equation 1) |  |  | Volume Model (Equation 2) |  |  |
| --- | --- | --- | --- | --- | --- | --- |
| | n | $\gamma$ | Error | n | $\gamma$ | Error |
| Dendrites and Soma ROI | 1.71 | 8.125 | 56.99 | 1.87 | 0.097 | 39.07 |
| Axon ROI | 1.33 | 0.346 | 24.29 | 1.66 | 0.015 | 42.19 |

**Supplementary Figure 1:**
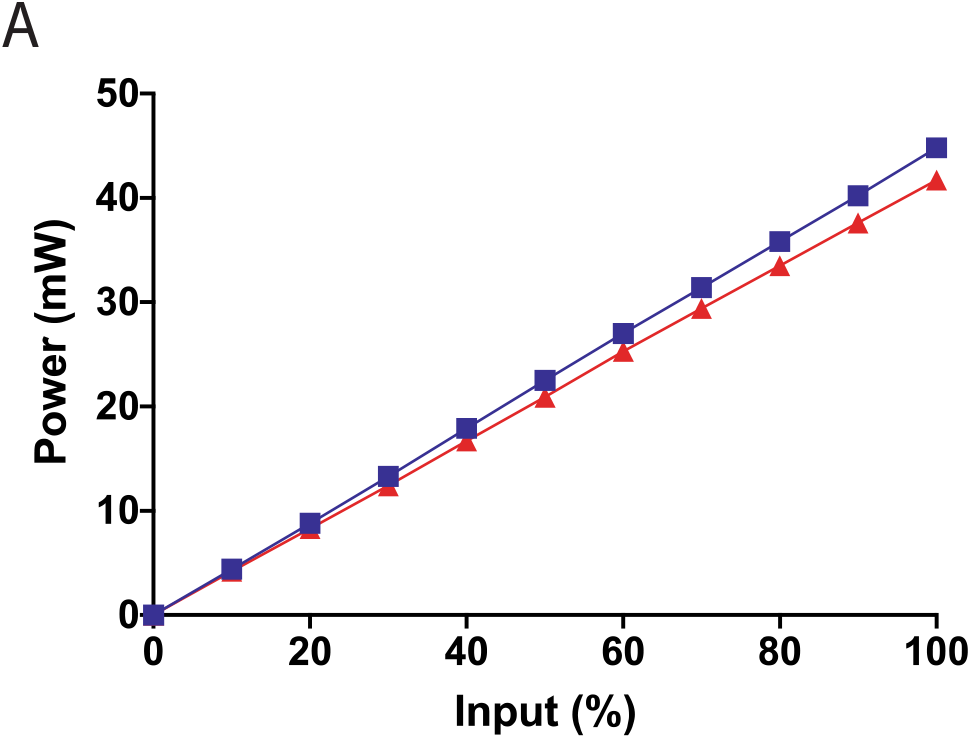
Measured power output (mW) across various input values (%) from 405-nm stimulus used to pulse Class IV neurons. Blue corresponds to stimulus with FWHM = 0.5 μm, red corresponds to stimulus with FWHM = 1 μm. Measurements were made using a microscope slide power sensor (S170C, Thor Labs) and a Touchscreen Optical Power and Energy Meter Console (PM400, Thor Labs) at the sample plane.

**Supplementary Figure 2:**
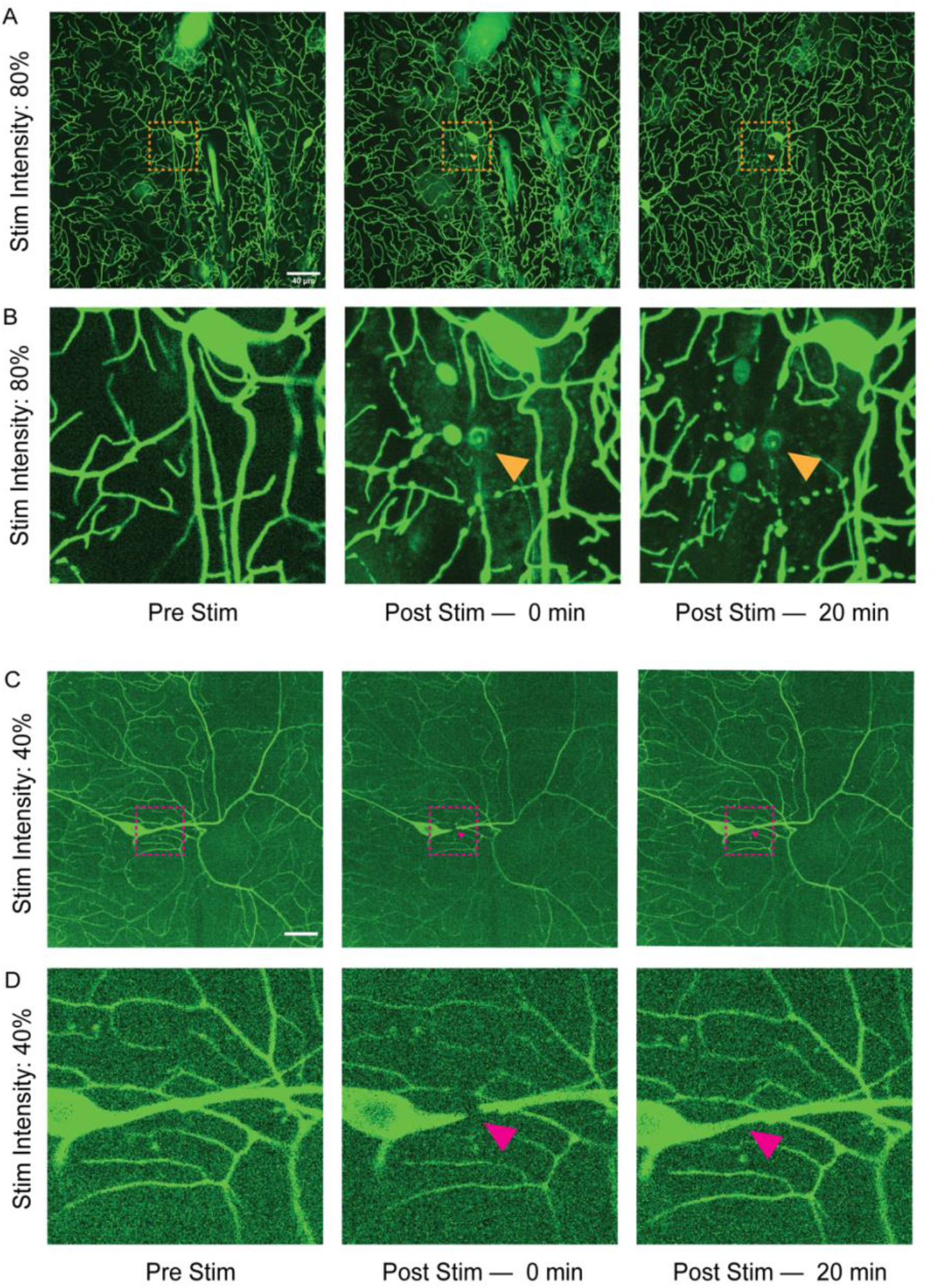
405-nm stimulation causes bleaching versus puncture of larval cuticle depending on the wattage delivered. Panels shown are pre-stim (left), frame immediately following stimulation (center), and 20 minutes post-stimulation (right). (A) Montage of Class IV da neurons expressing CD4-td-GFP stimulated with 405-nm laser at 80% power. (B) Zoom-in of region indicated by the dashed magenta box shown in panel (A). Orange arrow highlights region where laser was focused. Central and right panels show puncture wound that does not recover (not photobleaching) 20 minutes post stimulation. (C) Montage of Class IV da neurons expressing CD4-td-GFP stimulated with 40-nm laser at 40% power. (D) Zoom-in of region indicated by the dashed magenta box shown in (C). Magenta arrow indicates region where laser was focused. Central panel shows localized bleaching of dendritic process; right panel shows same region after recovery (20 minutes post stimulation) with no puncture wound.

**Supplementary Figure 3:**
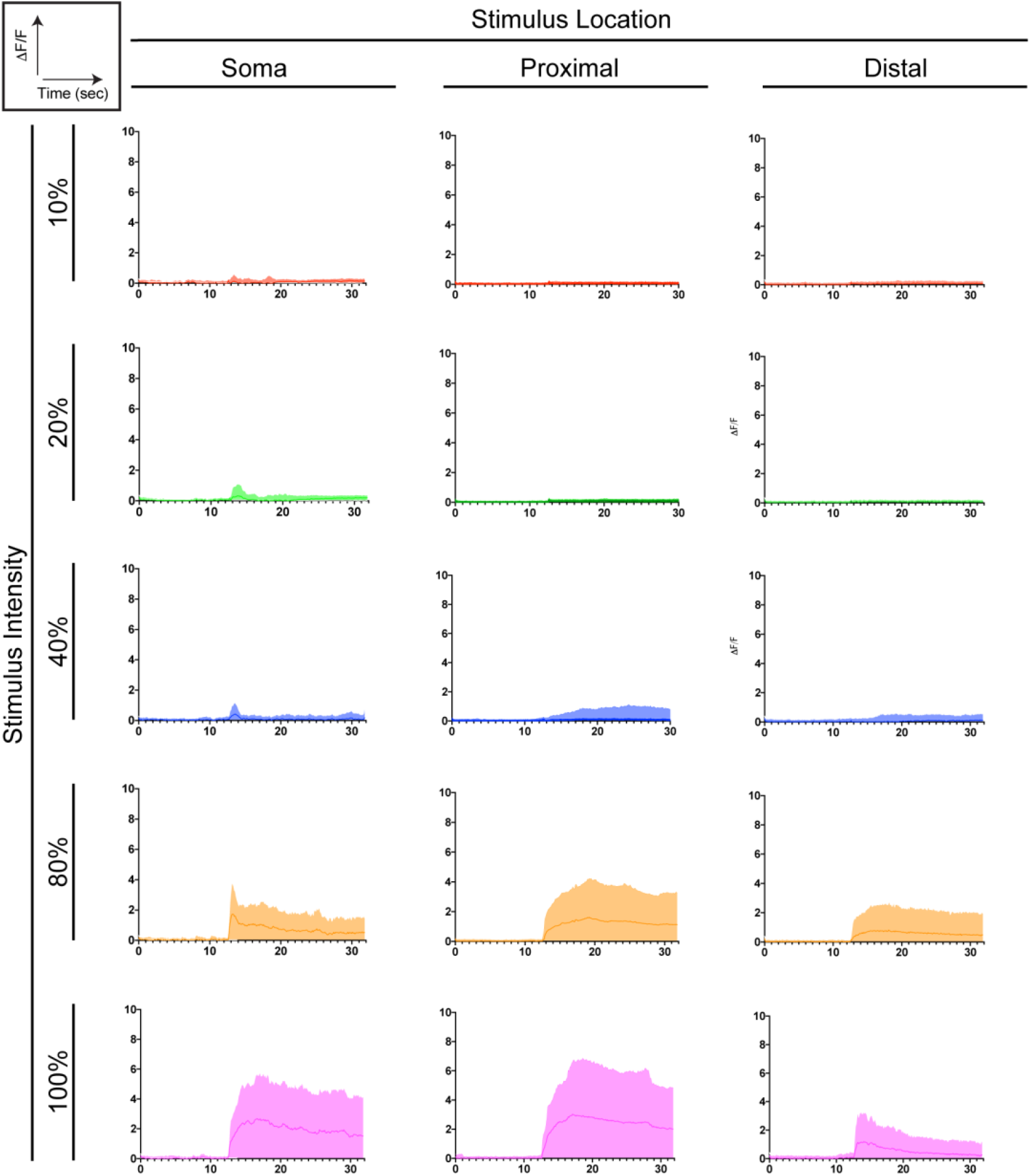
Representative *ΔF*/*F* traces of dendrite and soma ROIs for varying stimulation locations and wattages. Stimulus was activated at the 12.43 second mark (100th frame). Data across all experiments were combined. Mean and SD shown.

**Supplementary Figure 4:**
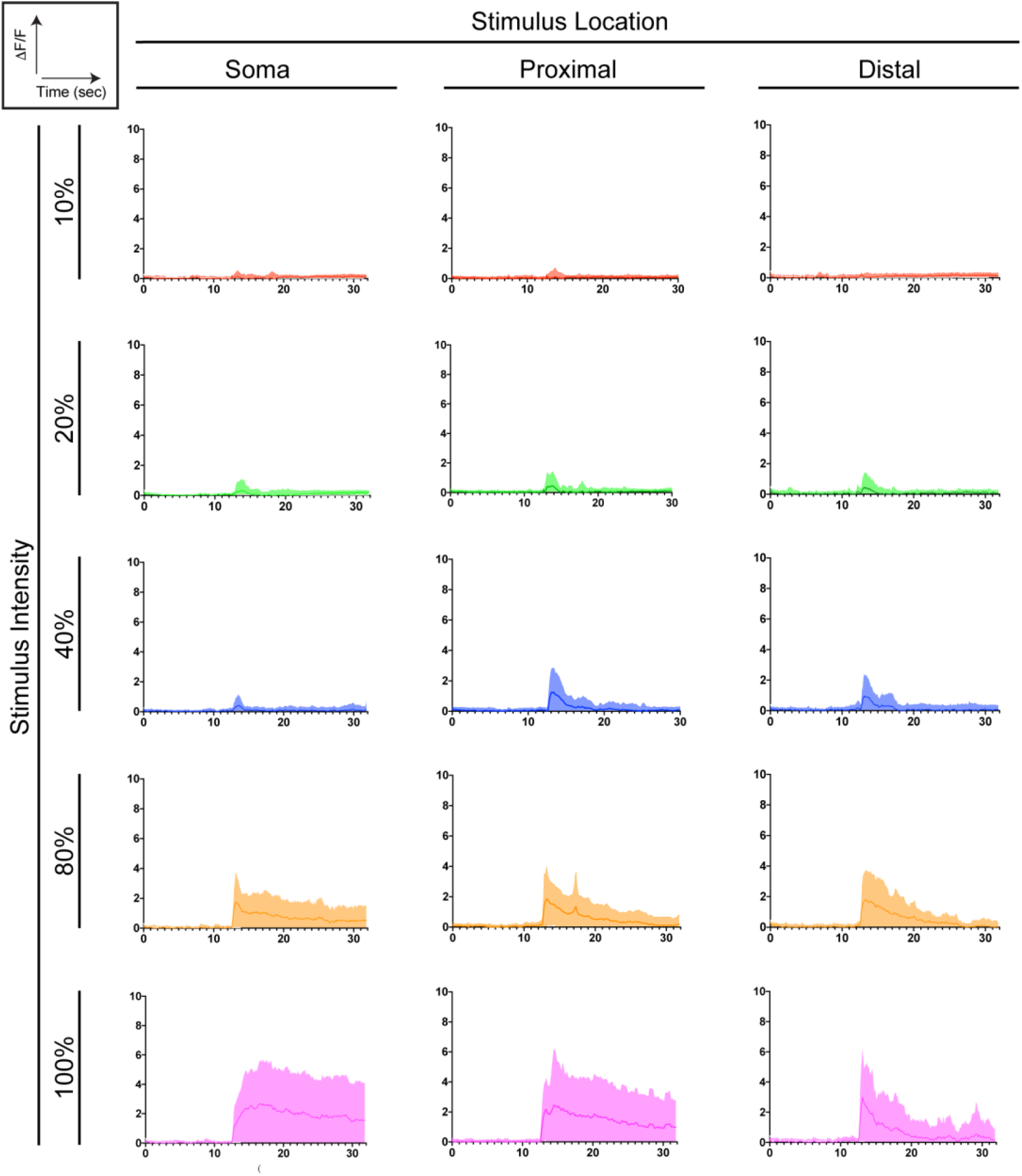
Representative *ΔF*/*F* traces of axon ROIs for varying stimulation locations and wattages. Stimulus was activated at the 12.43 second mark (100th frame). Data across all experiments were combined. Mean and SD shown

**Supplementary Figure 5:**
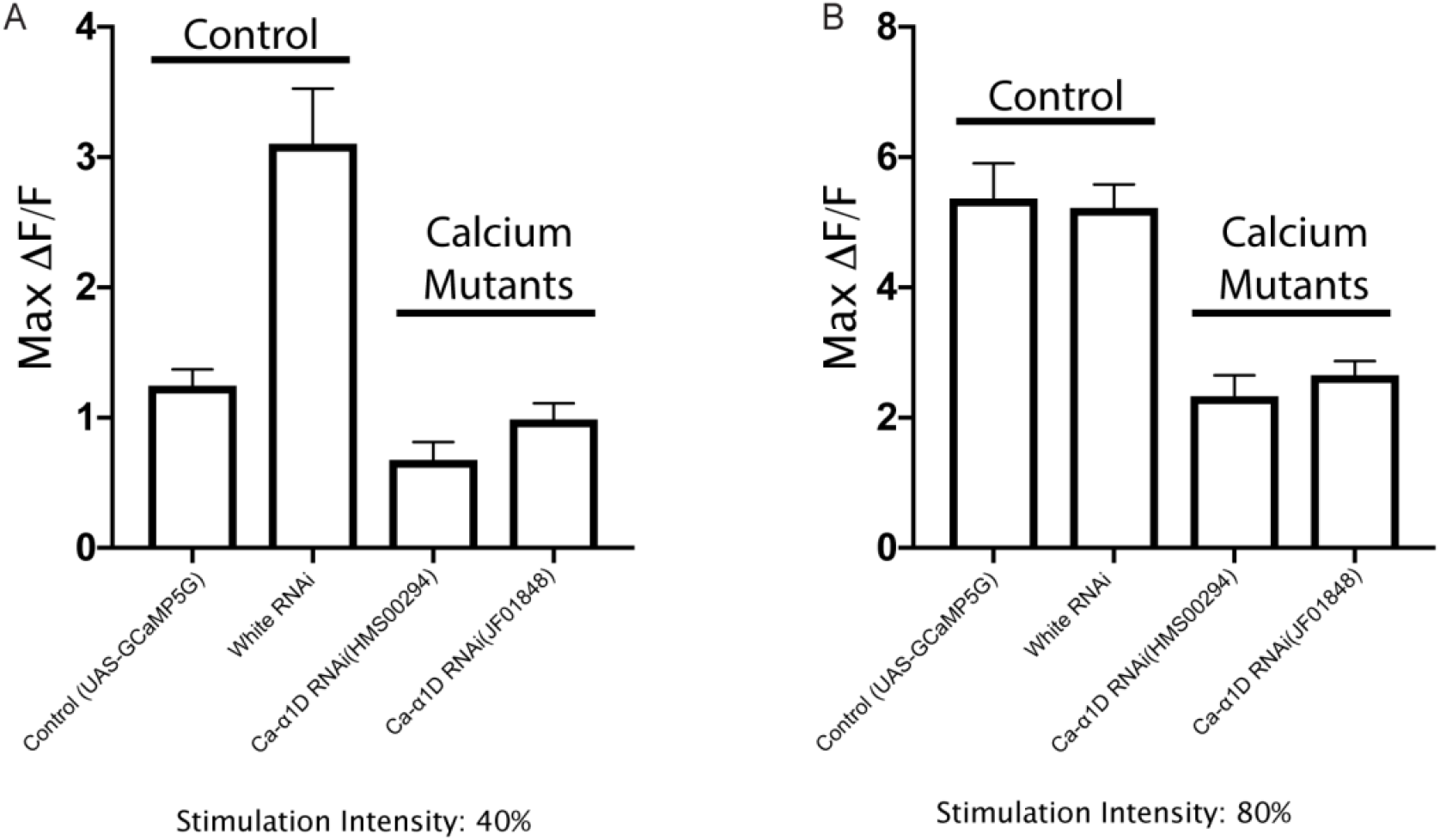
Magnitudes of calcium transients from control (; UAS-GCaMP5G / ppk-GAL4; | N = 24, n = 72 and; ppk-GAL4/+; white RNAi / UAS-GCaMP5G | N = 24, n = 72) and mutant cells (; ppk-GAL4/ +; *Ca-α1D RNAi* ^JF01848^ / UAS-GCaMP5G | N = 24, n = 72 and; ppk-GAL4/+; *Ca-α1D RNAi* ^HMS00294^/ UAS-GCaMP5G | N = 24, n = 72) irradiated at two different stimulation intensities, 40% (A) and 80% (B). Plots show mean and SEM.

### Links to Supplementary Movies

All Movies show Class IV da neuron expressing UAS-GCaMP6f. White dot indicates location of the laser stimulus.

**Supplementary Movie 1:** Behavioral Response at 80% Stimulation (Example 1) https://www.dropbox.com/s/5o8t1i3xwvtlesc/Supplementary%20Mov%201%20-%20BEHAVIORAL_RESPONSE_1.avi?dl=0

**Supplementary Movie 2:** Behavioral Response at 80% Stimulation (Example 2) https://www.dropbox.com/s/t0040zhmlswmjxm/Supplementary%20Mov%202%20-%20BEHAVIORAL_RESPONSE_2.avi?dl=0

**Supplementary Movie 3:** Non-contact axon response https://www.dropbox.com/s/gp6hw19vl7glat5/Supplementary%20Mov%203%20-%20NonContact_Axon_Response_80%25_Stim.avi?dl=0

**Supplementary Movie 4:** Contact dendrite and axon response https://www.dropbox.com/s/rm1d4nnaa7df962/Supplementary%20Mov%204%20-%20Contact_Dendrite_Response_80%25_Stim.avi?dl=0

**Supplementary Movie 5:** Contact Response (40% Stimulation Intensity) https://www.dropbox.com/s/1rtea8jqk9a0n4w/Supplementary%20Mov%205%20-%20ppkGAL4_GCamp6f_40%25_Stim.avi?dl=0

**Supplementary Movie 6:** Contact Response (80% Stimulation Intensity) https://www.dropbox.com/s/nfgpjmmde82zi2u/Supplementary%20Mov%206%20-%20ppkGAL4_GCamp6f_80%25_Stim.avi?dl=0

